# Generating brain-wide connectome using synthetic axonal morphologies

**DOI:** 10.1101/2024.10.04.616605

**Authors:** Remy Petkantchin, Adrien Berchet, Hanchuan Peng, Henry Markram, Lida Kanari

**Affiliations:** Blue Brain Project, EPFL, Chemin des mines 9, Geneva, 1202, Switzerland; New Cornerstone Science Laboratory, Institute for Brain and Intelligence, Southeast University, Nanjing, China; Shanghai Academy of Natural Sciences, Shanghai, China

**Keywords:** computational synthesis, axonal projections, brain connectivity, data-driven clustering, artificial neural network

## Abstract

Recent experimental advancements, including electron microscopy reconstructions, have produced detailed connectivity data for local brain regions. On the other hand, for inter-regional connectivity, large-scale imaging techniques such as MRI are best suited to provide insights. However, understanding the relationship between local and long-range connectivity is essential for studying both healthy and pathological conditions of the brain. Leveraging a novel dataset of whole-brain axonal reconstructions, we present a technique to predict whole-brain connectivity at single cell level by generating detailed whole-brain axonal morphologies from sparse experimental data. The computationally generated axons accurately reproduce the local and global morphological properties of experimental reconstructions. Furthermore, the computationally synthesized axons generate large-scale inter-regional connectivity, defining the projectome and the connectome of the brain, thereby enabling the *in silico* experimentation of large brain regions.

## 1 Introduction

Computational simulation and synthesis of the brain are essential to study the structural and functional complexity of the brain [1]. To this end, various methods have been developed for synthesizing dendritic neuronal morphologies and networks. Biophysically accurate models simulate neural growth by incorporating the molecular mechanisms of neuronal development [2], though their computational intensity often limits scalability. In contrast, phenomenological models, based on mathematical principles [3, 4] or statistical sampling from morphological datasets [5, 6], provide more computationally efficient alternatives. However, these models frequently require manual adjustments and struggle to generalize across different datasets. In 2022, we introduced a computationally efficient method for synthesizing biologically realistic dendritic morphologies [7], based on the topological morphology descriptor [8, 9]. A similar approach was employed to synthesize astrocytes within the complex neuro-glia-vasculature network [10]. However, the majority of models focus on dendritic synthesis and do not account for brain topology when generating morphologies.

In this work, we address the critical limitation of previous methods in synthesizing axonal trees. Unlike dendrites, which are generally restricted within a brain region, despite their environmental sensitivity [11–13], axons typically extend to specific brain regions and span much larger scales [14]. This highlights the necessity of generating axons that are spatially embedded within the brain [15]. Although there have been modeling efforts in areas such as axon guidance [16], trade-offs between material and conduction delay [17], and fiber arrangement [18], to date only the ROOTS model [19] has tackled the challenge of synthesizing detailed axonal morphologies. However, the axons generated by this latter model are restricted to the rat dentate gyrus and do not extend across brain regions, limiting their broader applicability to long-range axons.

Brain-wide long-range axons (LRAs) are the primary drivers of inter-regional connectivity [20] and plasticity [21]. Synthesizing LRAs presents a significant challenge, as it requires replicating the projection patterns and morphometric features of biological LRAs while also considering brain topology. In addition, biological data on LRAs was historically limited to just a few dozen morphologies until recent experimental advances in imaging and reconstruction techniques that enabled the reconstruction of hundreds of axons [22]. Successfully synthesizing brain-wide axons allows for the creation of synthetic connections between cells across the brain, thereby progressing from the connectome, encoding connections between individual neurons, to the projectome [23], which encodes connections between regions. Once this connectivity is established, it becomes possible to expand local simulations [24] to simulation of circuits involving multiple brain regions, getting closer to realistic full brain simulations.

This work introduces the first complete pipeline for synthesizing biologically realistic axons using sparse biological data. To generalize axon targeting from the available biological inputs, we employ an unsupervised Gaussian Mixture Model (GMM) clustering approach to group the input morphologies based on their projection patterns. This step ensures that the biological input projection patterns and proportions are reproduced. The resulting clusters serve as inputs for a long-range axon synthesis algorithm we developed [25]. The synthesized axons then traverse the brain, reaching their target regions according to their cluster assignments, with a preference to follow the path through fiber tracts to generate more realistic axonal shapes.

This species-agnostic workflow is applied to a dataset of brain-wide axons from the mouse brain containing previously published [26, 27] and novel data (with same methods as [27]), producing the first simulation-ready synthetic mouse brain with long-range axons. In this work, we demonstrate our algorithms on excitatory cells of the isocortex, specifically emphasizing layer five of the primary motor area due to the larger data availability in this region. Long-range axons of pyramidal cells are computationally synthesized, while local axons are grafted onto interneurons, following the approach used in our previous work [7, 24]. We validate the synthesized axons by comparing their morphological, topological, and targeting properties, demonstrating statistical similarity to experimental reconstructions. We furthermore validate our results by comparing the proportions of targeting against experimental data [28] that were not used as input to test the emergent properties of our algorithm. Finally, connectivity, computed with a method based on spatial proximity (see methods 4.4), emerges statistically similar to the input biological axons, which concludes the validation.

## 2 Results

### 2.1 Clustering

We show in Fig. 1 an example of clustering of 59 axonal morphologies from the Mouse-Light dataset [26] that originate from the presubiculum region of the mouse brain. In figures 1A, B, and C, each column represents an axon, and each row is a region where it terminates. Axons are grouped into the GMM clusters (top row) and based on their parent brain area (left column). We imposed a number of five clusters for these axons to echo Wheeler et al. [29], who found five clusters using unsupervised hierarchical clustering of axonal projections originating in the presubiculum. In Fig. 1A, we clustered the axons based on the axonal path lengths in the regions and sampled the lengths of the same number of axons from these clusters in Fig. 1B. These sampled axons are *virtual* in the sense that we did not synthesize them; we only sampled their lengths to verify the clustering. Some noise can be observed because a scalar variance was selected for each cluster by the Bayesian Information Criterion (BIC) optimization. Nevertheless, the main projections could be reproduced for each cluster. In Fig. 1C, we used the number of terminals in regions as clustering feature. The clusters are defined slightly differently due to the unsupervised nature of the clustering. However, we show in Fig. 1D that both features can be used equivalently for clustering. In the latter figure, each point corresponds to a target region, for which we computed the sum of axonal path length versus the number of terminals, for all axons of [26]. A linear relationship was found, showing that one feature is proportional to the other.

**Fig. 1.**
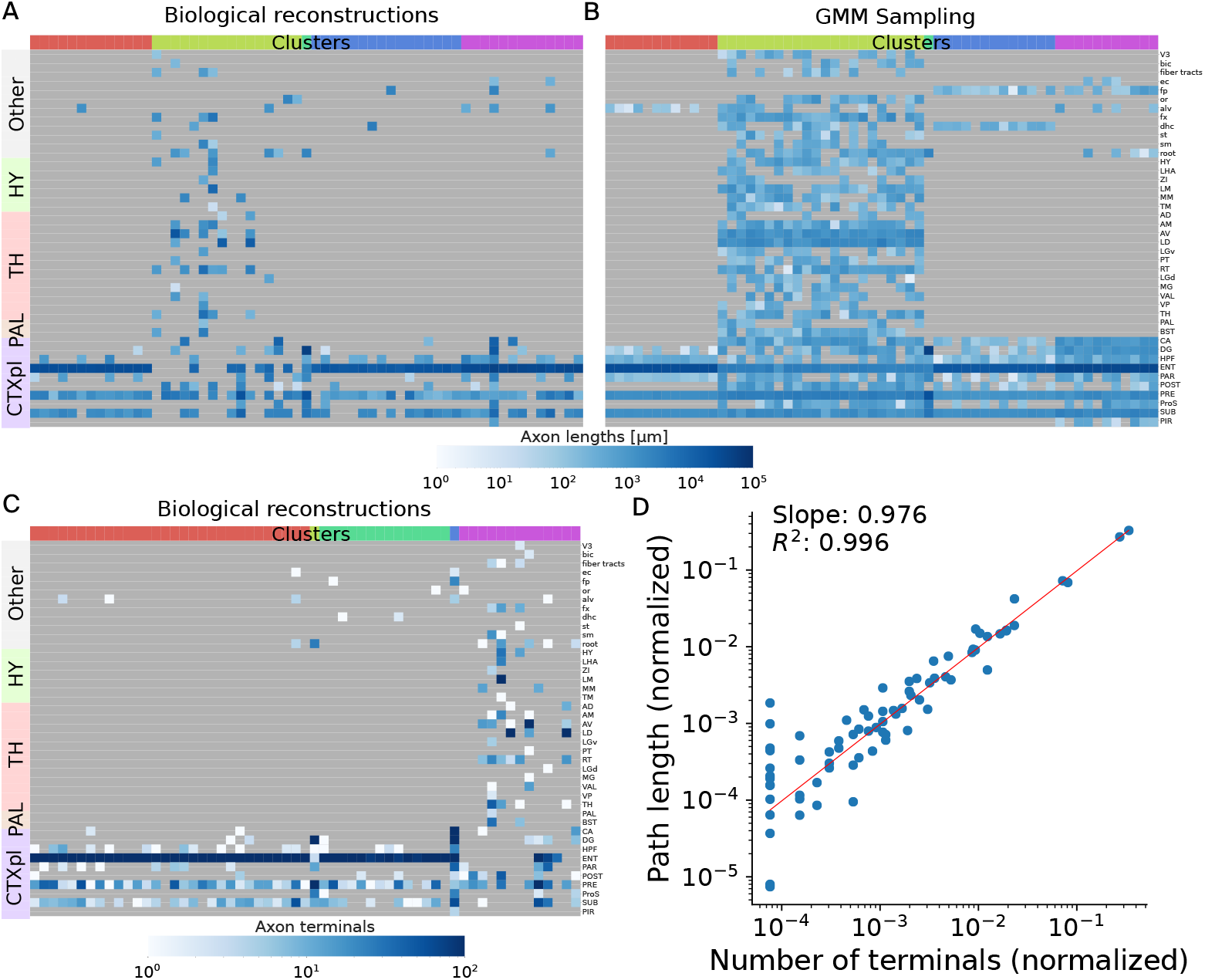
Clustering of 59 axonal morphologies from [26] with somata in the presubiculum. A total number of five clusters is imposed here because [29] reported five clusters in their clustering. **A**: Clustering based on axon lengths in target regions where the axons terminate. **B**: Projections of 59 virtual axons sampled with the GMMs defined by axon lengths (A). **C**: Clustering based on the number of terminals in the regions where the axons terminate. **D**: Using the number of terminals or axon lengths in target regions as feature vector for clustering was found to be equivalent. A linear relationship was found between number of terminals and path lengths of axons in the regions.

### 2.2 Synthesis in the mouse brain

In this section, we showcase two cases of LRA synthesis: the first originating from the primary motor area layer 5 (MOp5) and the second from multiple regions in the isocortex. We study the former in detail and use the latter as a window into the possibilities of this novel methodology. In both cases, we synthesized 54401 cells from the isocortex (approximately 1% of the actual density). Cells for which LRAs were not synthesized have grafted local axons as done in [7]. Local axons typically have a total path length of about 20 mm, whereas LRAs can go above 50 mm and up to 400 mm (distributions of lengths in supplementary Fig. S4B).

#### 2.2.1 MOp5 pyramidal cells

From the 54401 synthesized cells of the isocortex, 1695 were pyramidal cells from the MOp5, for which we synthesized LRAs in this section.

In Fig. 2, we offer a visualization of the 65 biological morphologies from the dataset that originated in the MOp5 (in blue), and 65 randomly picked synthesized morphologies with LRAs (in red). In Fig. 2A, we projected the reconstructions and the synthesized morphologies on a flat map of the isocortex, as described in [30]. In Figs. 2B and C, we used the `Brayns` tool [31] to visualize the mouse brain in three dimensions. We qualitatively see that synthesized axons follow the same paths and terminate in the same regions as the biological reconstructions. We proceed with a quantitative comparison of the two populations.

**Fig. 2.**
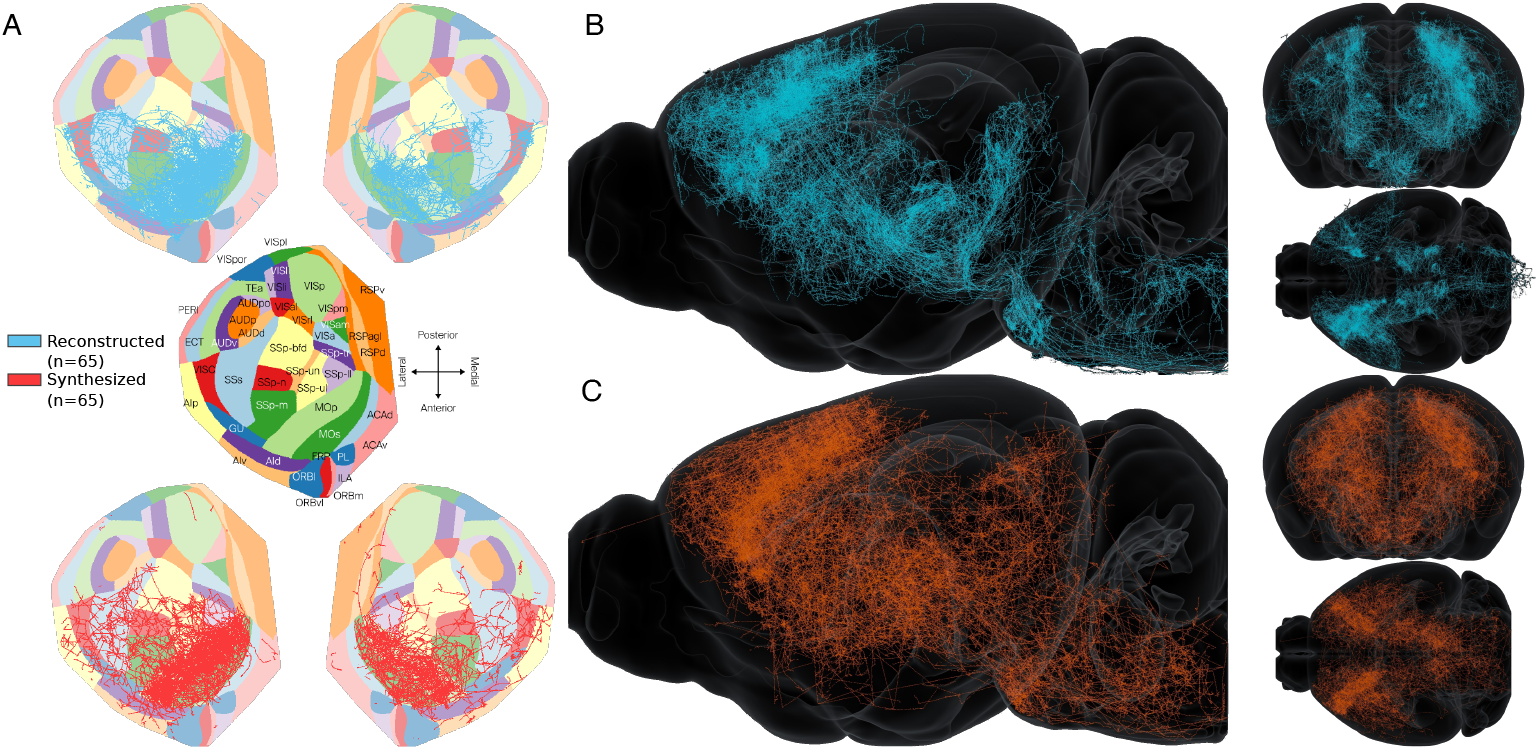
Comparison of 65 reconstructed (blue) and 65 of the 1695 synthesized axons (red) of MOp5 pyramidal cells. **A**: Flat map projection in the isocortex of the reconstructed and 65 randomly selected synthesized axons. **B**: The 65 reconstructed axons in the mouse brain atlas, in lateral (left), frontal and top views (right). **C**: 65 of the synthesized axons in the mouse brain atlas, lateral (left), frontal and top views (right).

##### Morphometrics

In Figs. 3A and B, we show a set of morphometrical features for the tufts (Fig. 3A) and trunks (Fig. 3B). Since there were more biological axons in the left (46) versus right (19) hemisphere, for the subsequent comparisons, we considered only a subset (362) of synthesized axons from the right hemisphere, to keep the same proportions. The distributions were normalized as in [7], where the value for an axon *a* was centered to the median of the reconstructed population 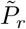, and divided by the standard deviation of the reconstructed population *σ*(*P*_*r*_):

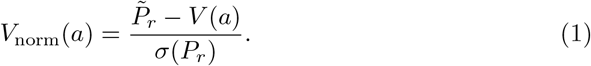

**Fig. 3.**
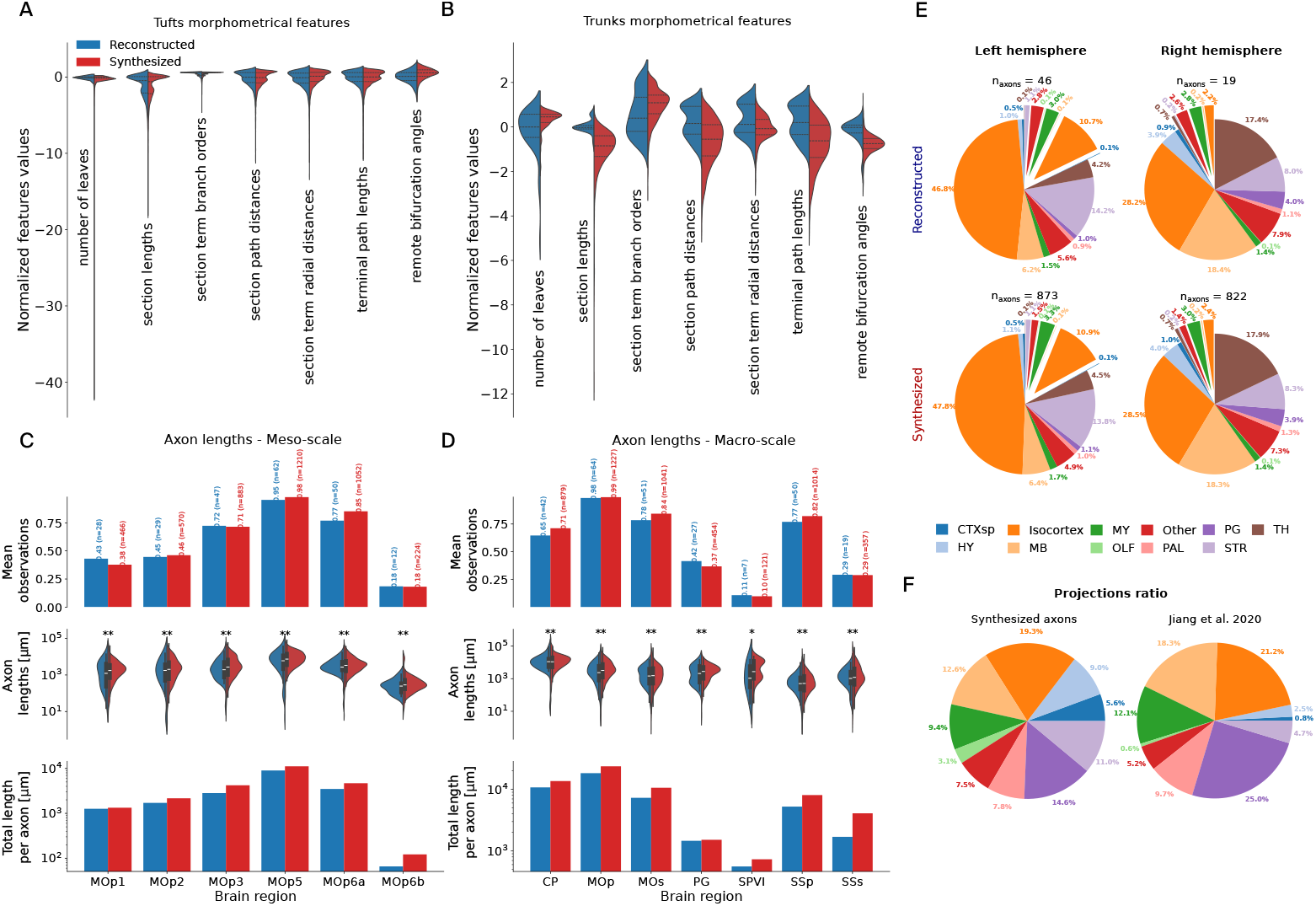
**A, B**: Morphometrical feature distribution comparison between the tufts (A) and trunks (B) of the reconstructed (blue) and synthesized (red) axons. All distributions were centered to the reconstructions median and normalized by the reconstructions standard deviations, as done in [7], see equation (1). Medians, first and third quartiles are shown (hyphens, dotted lines). **C, D**: Comparison of axonal lengths in regions of interest between reconstructed and synthesized axons. Top: proportion of axons terminating in the regions. Middle: distribution of axon lengths. Medians, first and third quartiles are shown in the inner boxes. Bottom: total length normalized by the number of axons. Fig. C focuses on the MOp layers and D on distal regions. **E**: Comparison of tufts locations clustered from the reconstructed (top) and synthesized (bottom) MOp5 axons. The exploded parts in the pie charts represent projections in contralateral regions. **F**: Projections ratio comparison between our synthesized axons and reconstructed MOp5 axons analyzed in [28].

The synthesized and clustered tufts morphometrical features are in good agreement. The tufts were synthesized with the dendrites synthesis algorithm, which has already proved to reproduce well morphometrics of input dendritic trees [7]. This shows that dendrites and tufts can be mathematically described with the same topological method. Morphometrics of the axon trunks were found to match reasonably well with biological inputs when they were mimicked, as in Fig. 6 and [25]. However, with the new selection of target points presented in this work, many morphometrics, such as the number of leaves, angles, and those related to distances and lengths, are subject to change. This resulted in overall longer section lengths and path distances, slightly increased remote bifurcation angles, and slightly fewer leaves and branches. We finally compared the path lengths of the axons in targeted regions in Figs. 3C and D. Fig. 3C focuses on the layers of the MOp (meso-scale), and Fig. 3D on the set of regions highlighted in Fig. S3 (macro-scale): the caudoputamen (CP), the primary and secondary motor areas (MOp, MOs), the pontine gray region (PG), the spinal nucleus of the trigeminal interpolar part (SPVI), and the primary and secondary somatosensory areas (SSp, SSs). The top subplot of Figs. 3C and D shows the proportions of reconstructed (blue) and synthesized (red) axons terminating in the regions; in the middle subplot, the distribution of axon lengths, and finally, in the bottom subplot, the total lengths of all axons, normalized by the number of axons in the populations. The stars in the middle subplots show strong significance (**) and non-randomness (*) when the Maximum Visible Spread (*MVS*) score between the populations is < 0.1, respectively < 0.5 (see section 4.2 and [7] for a definition of the MVS). Overall, we can see that the terminating number of axons and their path lengths in the targets were accurately replicated.

##### Targeting

We now compare the location of clustered and synthesized tufts in Fig. 3E, by looking at the location of the common ancestors of the tufts computed in the reconstructed axons (top row) and placed in the synthesized axons (bottom row). We only show a set of brain regions, which were reported to be targeted by 42 MOp5 axons in the dataset analyzed in [28]: the cortical subplate (CTXsp), isocortex, medulla (MY), PG, thalamus (TH), hypothalamus (HY), midbrain (MB), olfactory areas (OLF), pallidum (PAL), and striatum (STR). The exploded slices of the pie charts show targets in the contralateral hemisphere, whereas the normal slices are in the ipsilateral hemisphere. We see that the proportions of targets clustered and synthesized and the number of axons starting from each hemisphere were statistically well reproduced. In Fig. 3F, we compared the projection ratio from a different set of biological reconstructions from [28] and our synthesized axons. The projections ratio is defined as the proportion of axons targeting each region. Even though we used a different source for biological inputs in this work, the synthesized axons primarily targeted the same areas, and their projections ratio was not far from [28], with the highest difference of 10.4% in PG, and around 5 to 7% in CTXsp, HY, STR and MB.

##### Connectivity

Finally, we computed the connectivity of the MOp5 pyramid cells and show the results in Fig. 4. We chose to display here the connections in all the subregions of the isocortex: the anterior cingulate area (ACA), frontal pole of the cerebral cortex (FRP), gustatory areas (GU), infralimbic area (ILA), MOp, MOs, orbital area (ORB), prelimbic area (PL), retrosplenial area (RSP), SSp, SSs, auditory areas (AUD), ectorhinal area (ECT), perirhinal area (PERI), temporal association areas (TEa), visceral area (VISC), visual areas (VIS), agranular insular area (AI) and posterior parietal association areas (PTLp).

**Fig. 4.**
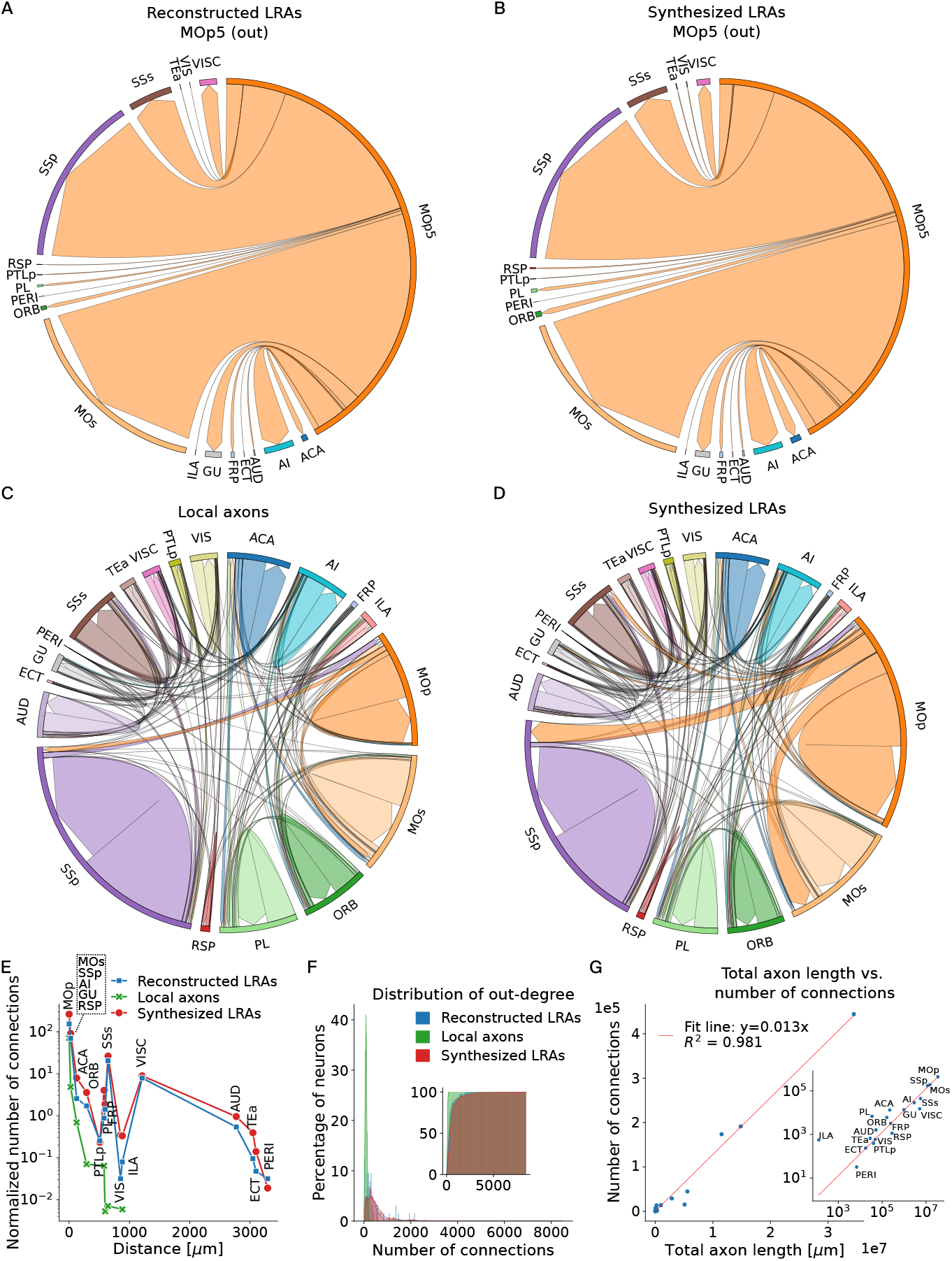
Connectivity of the synthesized MOp5 axons. **A, B**: Proportions of outgoing connections from MOp5 pyramidal cells, using the 65 biological reconstructions (A) and the synthesized LRAs (B). **C, D**: Projectome of the isocortex subregions, with local axons only (C), respectively, with long-range axons for the MOp5 pyramidal cells (D). **E**: Number of connections in subregions of the isocortex versus distance to subregion, for biological LRAs (blue), local axons (green) and LRAs synthesized with the present method (red). As can be expected, local axons connect only to regions close to the MOp. **F**: Out-degree distribution of the biological LRAs (blue), local axons (green) and synthesized LRAs (red). Inset: cumulative histogram. **G**: Total number of connections vs. length of the synthesized long-range axons in the subregions of the isocortex. A linear relationship seemed to describe the observed data well.

First, we compared in Figs. 4A and B the proportion of outgoing connections of the 65 biological reconstructions from MOp5 (A) and the synthesized axons of the MOp5 pyramidal cells, with the same proportions in both hemispheres (B). The arrows represent the direction of efferent axo-dendritic synaptic connections, and their size is proportional to the number of connections. We did not show in this figure the connections within MOp because all the synthesized cells in the MOp5 region were removed when the reconstructions were used (Fig. 4A). This was to ensure that only the connections of the reconstructions were shown and not those of the local grafted axons, which would have biased the result. As can be seen from the figures, the connection proportions are statistically equivalent.

We then plotted the projectome of all the synthesized cells of the isocortex, without the LRAs (local axons only) in Fig. 4C and with the synthesized LRAs in Fig. 4D. The projectomes with only the outgoing connections can be seen in supplemetary Fig. S2. We can see that only by synthesizing LRAs for the MOp5 pyramidal cells has the projectome significantly changed.

In Fig. 4E, we computed the number of connections formed in the isocortex subregions by the MOp5 local axons, reconstructed and synthesized LRAs, and plotted them against the distance to each subregion. The distance to the subregions was computed as the closest distance between the border of the MOp region to the border of the other regions. We can see that, as expected, the local axons connect only to areas close to the MOp. Also, we see that the number of connections of synthesized LRAs reproduced the connections of the reconstructed LRAs well. Fig. 4F shows the distribution of out-degree of the local axons, reconstructed and synthesized LRAs. Not only the number of connections of long-range axons was much higher and happened in more regions, but also the shape of the distribution was different, as can be seen in the inset of Fig. 4F. Where the local axons out-degree seemed to follow an unimodal law with small number of connections, the distribution of LRAs seemed to have a long tail towards a high number of connections. The out-degree distribution of synthesized LRAs agrees with and generalizes the out-degree distribution of the reconstructed LRAs. We found that the number of connections was proportional to the total axon lengths in brain regions; see Fig. 4G. This can also be seen in the supplementary Fig. S4A, where the shape of the distribution of axons total lengths resembled the shape of the out-degree distribution of LRAs. This result could help predict the connectivity in brain regions. However, one should be aware that the coefficient of this relationship depends on the number of cells synthesized in the regions, up to a limit (such as the total number of boutons of the axons or the maximum possible number of dendrites in the region).

These results show that synthesizing LRAs is essential for connecting cells locally and across distal brain regions.

#### 2.2.2 Isocortex pyramidal cells

Finally, to showcase the possibilities of the present method, we synthesized LRAs for the pyramidal cells of all subregions of the isocortex, for which we have biological inputs to build GMM clusters. A total of 1472 biological axons were used as input for these regions: MOp (*n* = 159), MOs (*n* = 393), SSp (*n* = 488), SSs (*n* = 97), VISC (*n* = 8), VIS (*n* = 83), AUD (*n* = 26), ECT (*n* = 4), GU (*n* = 11), ORB (*n* = 24), ACA (*n* = 40), RSP (*n* = 46), FRP (*n* = 7), PL (*n* = 8), TEa (*n* = 10), PTLp (*n* = 26), AI (*n* = 42). A summarizing table showing the count per layer and hemisphere can be found in the supplementary material, table S2. This amounted to a total of 21680 synthesized long-range axons originating from the isocortex subregions. In Fig. 5A and B, we visualized the synthesized cells in the mouse brain. Cells are colored by the cortical layers of their somata in Fig. 5A and by region of origin in Fig. 5B. The region colors correspond to the color mapping in Fig. 5C. The projectome shown in Fig. 5C, although incomplete due to the sparsity of input data, shows a fundamentally different amount and pattern of connections than the projectome generated with only local axons in Fig. 4D. The motor and somatosensory areas showed much more efferent and afferent synaptic connections, and regions were much more interconnected overall.

**Fig. 5.**
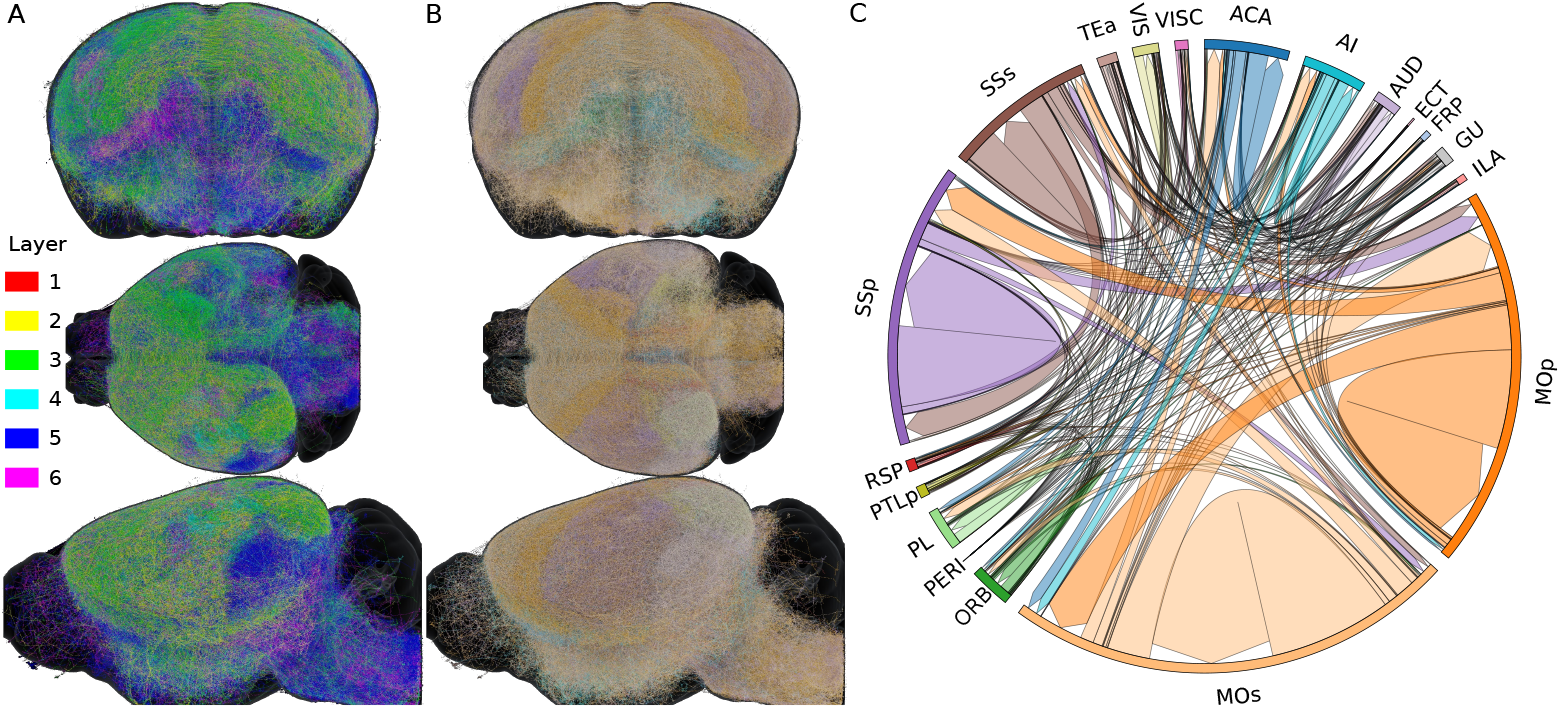
Synthesis of 21680 axons for pyramidal cells in subregions of the isocortex. The list of regions for which we had biological data to synthesize the axons can be found in the supplementary table S2. **A, B**: Visualizations in the mouse brain of the synthesized axons, colored by layer, respectively by region, of their somata. 25% of the synthesized axons are shown. The color code of Fig. B is the same as Fig. C. **C**: Projectome of the long-range axons. Arrows show the direction of connection (from preto post-synaptic), and their size is proportional to the number of connections.

## 3 Discussion

In this work, we presented a workflow for synthesizing simulation-ready, detailed longrange axonal morphologies of the mouse brain. While we applied this method to the mouse brain, it can be adapted for rat or human brains with minimal parameter tuning, depending on data availability. We demonstrated that the workflow successfully reproduced the proportions, lengths, and number of tufts in biological axons within targeted regions, with the tufts exhibiting statistically similar morphometrics. Connectivity within regions was accurately replicated, as shown in [7], where reproducing similar dendritic morphologies led to equivalent local connectivity between biological and synthesized circuits. Long-scale connectivity is also consistent with experimental data. In our validation instance of the MOp5 region, axons targeting was even consistent with morphologies not included in the algorithm inputs. Our algorithm is thus capable of generating full brain connectivity predictions, that can be compared against and completed from future experiments, once more data becomes available.

Small differences in axon lengths and resulting connections are likely due to the random placement of tuft target points within the target regions, occasionally causing tufts to extend beyond the intended areas. This issue could be mitigated by either constraining tuft growth within the target regions or positioning them closer to their common ancestor in the reconstructed dataset. In addition, as discussed in [25], improvements in the synthesis algorithm will improve the agreement between synthesized and biological axons. These include a refined version of the Steiner tree algorithm to incorporate more constraints, as well as an optimization process to select the most appropriate input parameters based on the input datasets. In addition, as more longrange axonal data become available, we will be able to better approximate biological rules, and improve the ability of our algorithm to generalize to new inputs.

Our approach is versatile, due to the small number of parameters, that require minimum tuning from the user. By focusing on replicating the biological data while allowing for statistical variability, we enable properties such as connectivity to emerge naturally, rather than imposing them onto the model. However, this dependence on biological data also introduces some limitations. We are constrained by the projection patterns present in the input dataset, thus we can only accurately synthesize axons for the entire brain if all relevant projection patterns are represented. Additionally, the targeting of the synthesized axons is sensitive to the quality of the reconstructed axons in the dataset. In the future, we will expand this methodology to generalize the observed patterns in the full brain, also linking the observed projection patterns to gene expression through transcriptomic datasets [32–34]. In addition, the clustering method could be improved by considering region hierarchy or spatial location within regions in the clustering step. Finally, we could leverage the Gaussian nature of the clusters to sample the lengths in the synthesis step.

One of the key advantages of our algorithm is its ability to synthesize wholebrain axons, leading to a comprehensive model of full-brain connectivity. Therefore, we enable the computational synthesis of a full brain, once a more complete axonal projection dataset becomes available. By integrating additional biological data, we will refine and extend our models to simulate more complex and realistic brain-wide connectivity. The scalability and adaptability of our method make it a valuable tool for future research, bridging the gap between current experimental limitations and the goal of creating a fully synthetic, computational model of the brain for computational simulations [24, 35].

Looking further into the future, our methodology holds significant promise for advancing healthcare applications. The generation of highly accurate and biologically informed neural circuits can significantly enhance patient diagnostics, particularly for conditions involving abnormal brain connectivity. For example, our algorithm will be useful for large-scale MRI simulations [36] to improve neuro-imaging predictions for improving patient diagnostics. This approach could allow for earlier detection of neurological disorders by simulating disease progression and identifying biomarkers before symptoms manifest. Additionally, the method’s scalability and ability to synthesize diverse axonal projections can contribute to more efficient drug discovery by facilitating *in silico* testing of therapeutic compounds across realistic brain networks. Furthermore, the model’s adaptability to individual datasets makes it well-suited for personalized medicine by facilitating the simulation of patient-specific brain connectivity and drug responses, which could revolutionize treatment strategies and optimize long-term outcomes for patients.

## 4 Material and Methods

### 4.1 Nomenclature

In this work, we used the terms “brain region” in a broad sense: it might label entire regions (thalamus, cerebellum), sub-regions (primary, secondary motor areas), or layers separation (primary motor area, layer 5, layer 6).

We used brain regions acronyms defined in the Allen Brain Atlas [37]. We spelled out each acronym that is helpful for this work’s comprehension when they were used. We also provided an exhaustive list of all brain acronyms used in the supplementary section 7.1.

### 4.2 Long-range axon synthesis

We presented a novel algorithm for synthesizing long-range axons (LRAs) in [25]. This novel method was able to synthesize cells accurately mimicking reconstructed axonal morphologies. We review here the main relevant steps of this algorithm; see Fig. 6A. Reconstructed axonal morphologies are taken as input. Terminations that are close together (within a radial and path distance) are clustered up to their common ancestors in so-called *tufts*. The axons with all tufts taken out are labeled as *trunks*. The somata of the reconstructed morphologies are used as *source points*, and the locations of the tufts ancestors as *target points*. A graph is created with these and additional points taken as vertices (see [25] for details). Then, we run the weighted Steiner tree algorithm [38] on this graph, which connects the targets while minimizing cable length, and allowing to prefer edges, such as ones that would be inside fiber tracts. The created trunk is then post-processed to reproduce local morphometrics of biological axons, with operations such as adding random noise and taking into account local history for modifying the curvature. Finally, the tufts that were initially clustered are synthesized based on their topological properties, with the method previously described and validated for synthesizing dendritic morphologies [7].

**Fig. 6.**
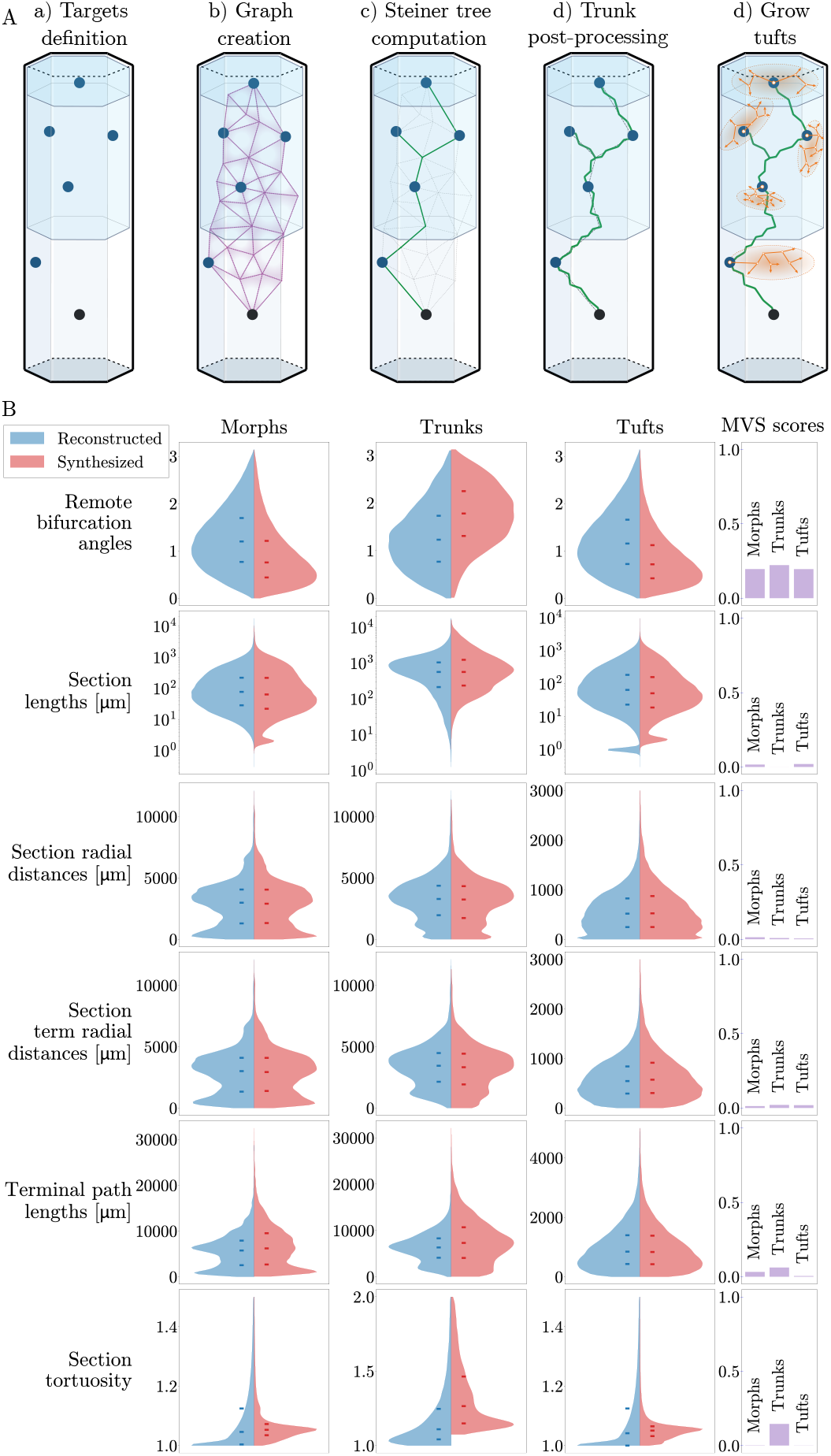
**A**: Main steps of the long-range axon synthesis. From an input biological axon, we take the soma as the source point, cluster the tufts, and make their common ancestors the target points. The synthetic trunk is created by connecting the targets from the source point with the Steiner tree on a graph method. The trunk is then post-processed with the addition of noise to reproduce local morphometrics of biological axons. Finally, the tufts are grown at the target points from their topological properties, with the same method as dendrites [7]. **B**: Distribution of morphological features across 1076 reconstructed (blue) and synthesized (red) axonal trunks (second column), tufts (third row), and complete morphologies (first column). The medians, first and third quartiles are shown on the distributions (hyphens). The MVS score is plotted for each morphometric distribution in the last column.

We could assess that this algorithm was able to produce morphometrical properties that matched with the reconstructed axons at the trunk, tuft, and morphology levels, see Fig. 6B. We measured a maximum value of 0.2 for the Maximum Visible Spread (MVS) score introduced in [7], which is a measure of the difference between the medians of two distributions with respect to their joint dispersion.

### 4.3 Axonal projections analysis

The formalism introduced in the axon synthesis algorithm takes all its meaning when the method which we present in this work is used along. In our previous work [25], source points and target points were directly taken as the somata and common ancestors of tufts of the reconstructed morphologies, respectively. Furthermore, the topological signature of the tufts of the biological axon to mimic was used for their synthesized analogs. In the present work, we generalized the selection of source points, target points, and tufts from populations of biological axons rather than from individual morphologies. This eventually allowed us to synthesize and connect neurons in the entire mouse brain, within and between brain regions.

We explain our methodology in what follows, see Fig. 7 for a visual schematic.

**Fig. 7.**
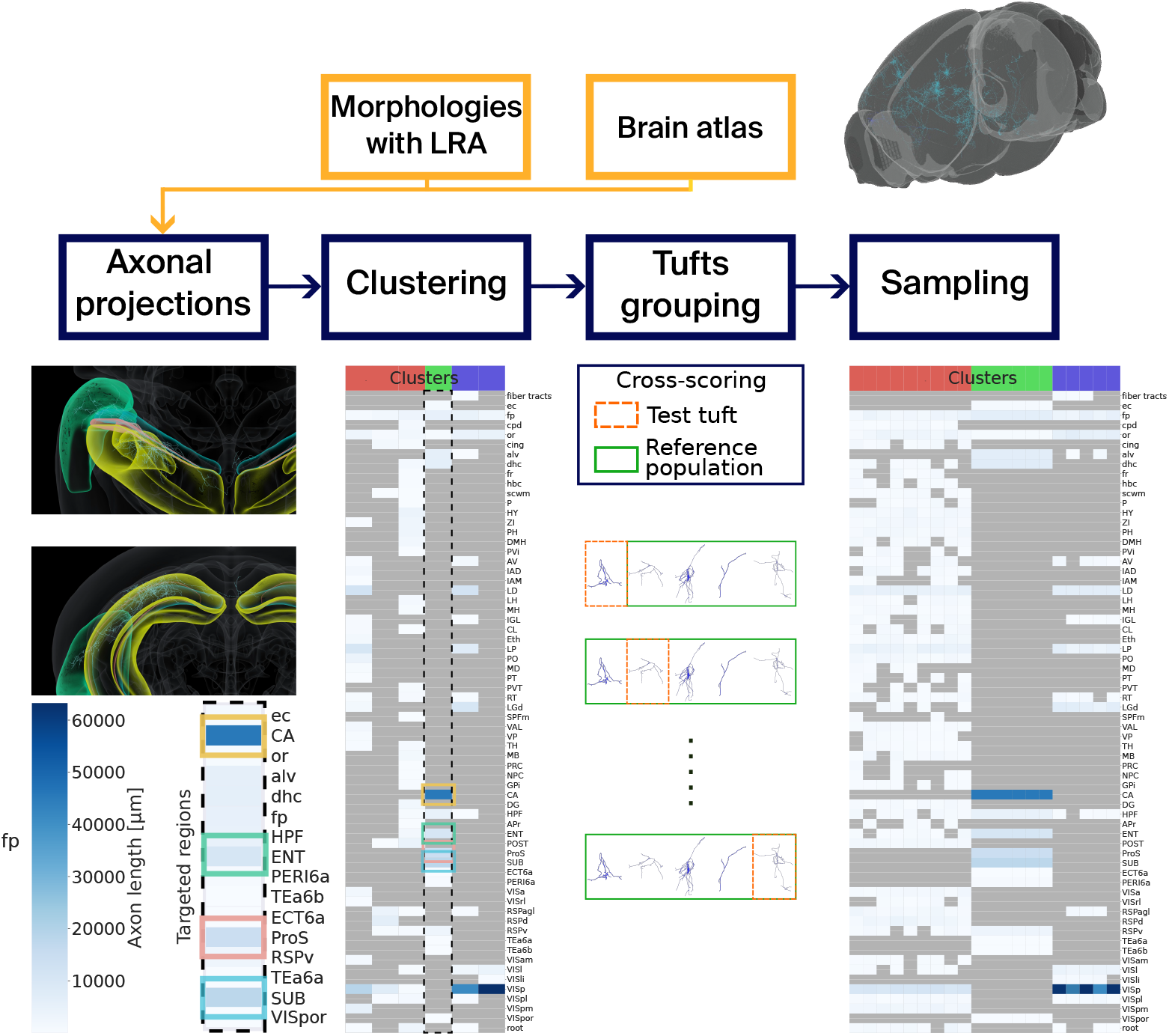
Schematic view of the axonal projections analysis. Neuronal morphologies with long-range axons (LRAs) placed in a mouse brain atlas are taken as input. The projections of the LRAs are computed, and we cluster the LRAs based on these projections with GMMs. Tufts are separated from the trunks of the LRAs, and they are counted and assigned a score for each cluster and target brain region. Finally, we can use the GMMs to sample target axon lengths for every target brain region defined in the clusters. fp: corpus callosum, posterior forceps.

#### 4.3.1 Input data

##### Morphologies

We used morphologies with long-range axons from three different sources:

- 1084 morphologies from the Janelia MouseLight Project [26]
- 1741 morphologies from [27]
- 800 novel morphologies from collaboration with Southeast University, Nanjing, China and H. Peng (same protocol as [27])

Morphologies could be incompatible with our methodology for reasons such as incomplete reconstructions or artifacts due to the reconstruction techniques. Therefore, we post-processed them with the `Repair` workflow of `Morphology-Workflows` software [39], which applied corrections such as repairing out-of-plane cut branches or removing unifurcations. 16 (8 from [26] + 8 from [27]) morphologies did not pass all the correcting processes, and 8 (1 + 5 + 2) did not pass the axonal projection analysis (due to being detected out of bounds or having faulty axons). In total, 3601 morphologies were used in this work.

##### Mouse brain atlas

We used the mouse brain atlas described in [30], which is an enhancement of the Common Coordinate Framework version 3 (CCFv3) atlas of the Allen Brain Institute [37]. Notable additions are the annotations of the barrel field areas and the distinction of cortical layers 2 and 3.

#### 4.3.2 Projections computation

In order to generalize the axon synthesis algorithm, we first computed the projections of reconstructed biological axons by counting the number of terminals of each axon in all regions they terminated and computing the axon path lengths in these regions. Our method then uses one of these features to cluster the axons with somata in the same source region.

Let us formalize this problem in the following way. We want to classify a set of *N* biological neurons. Let *a* be the axon of a neuron *n*. We denote *s*_*a*_ as the source brain region where the soma of neuron *n* is located. The axon *a* projects and terminates in a number of target brain regions, *t*_*a*_ ∈ (ℕ)^*B*^ is the vector counting the number of terminal points in all brain regions *B*. In other words, 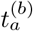 is the number of terminal points of axon *a* into brain region *b*. We define in a similar way *l*_*a*_ ∈ (ℝ^0+^)^*B*^ the path length of axon *a* inside brain region *b*. Note that brain regions can be defined at various levels of detail according to the hierarchy of the brain atlas.

Let us further denote *f*_*a*_, the feature vector for the classification of neuron *a*. We consider the case where *f*_*a*_ = *l*_*a*_. *s*_*a*_ is not included in *f*_*a*_ because we impose a separate classification for each source region.

#### 4.3.3 Clustering method

There are several methods for unsupervised clustering of data such as K-means and its variants (K-medoids, K-centroids [40]) and hierarchical clustering based on statistical difference of the subclasses [29]. We chose here to assume that our data could be described by normal probability density functions, using Gaussian Mixture Models (GMMs) [41].

Let us imagine that region *s*_*a*_ has *C* ∈ *ℕ* clusters. To assume that *a* originates from a GMM is equivalent to saying that the probability of *a* to belong to cluster *c* is given by :

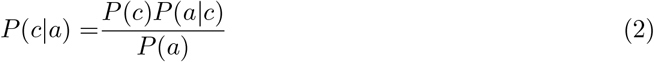

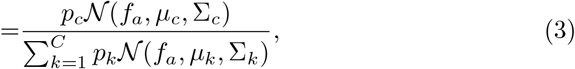

where 𝒩 (*f*_*a*_, *µ*_*c*_, Σ_*c*_) is the multivariate normal distribution with mean *µ*_*c*_ and covariance matrix Σ_*c*_:

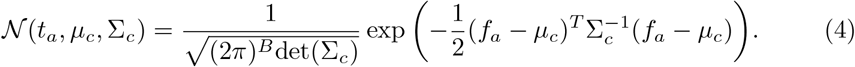

We assume Σ_*c*_ symmetric and positive definite ∀*c* ∈ *C*. Note that we have the constraint 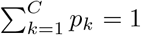.

A common technique [42] to optimize the clustering is to maximize the likelihood, or log-likelihood, of the observed data, based on the parameters of clustering *θ* [42]. Here, we can write *θ* = {*p*_1_, …, *p*_*C*_, *µ*_1_, …, *µ*_*C*_, Σ_1_, …, Σ_*C*_}. The log-likelihood of the data based on the parameters of clustering is given by (assuming *a* are iid) :

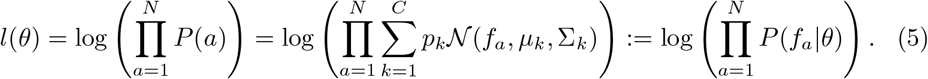

However, optimizing the log-likelihood (5) is not tractable in the case of large GMMs. Instead, we used the Expectation-Maximization (EM), see Supplementary material 7.2.1.

The number of clusters *C* for a source region can be either imposed or selected to maximize a score, the Bayesian Information Criterion (BIC), see Supplementary material 7.2.2. Since we were less interested in describing the biological data in this work than reproducing it, the number of clusters *C* per source region *s* is optimized on the BIC score within a range of *C* going from the number of axons *N*_*s*_ in *s* divided by 2, to *N*_*s*_, unless specified otherwise.

#### 4.3.4 Tufts grouping

Once the GMM clusters were defined for all source regions of the input axons, the tufts were clustered using the previously described clustering algorithm, section 4.2 and [25]. We used a maximum clustering radius and path distance of 300 *µ*m. The tufts were then grouped by GMM cluster and region of their common ancestors. For each group *g*, we computed the average 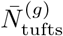 and variance 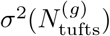 of tuft numbers for each group. Finally, the tufts were assigned a *representativity score* within their group, which is a measure of how close they are to the other tufts in their group in terms of a set of morphometrical features. In this work, all morphometrics were computed using `NeuroM` [43]. We used the MVS score to measure the similarity of the features. The calculation details can be found in supplementary material 7.3.

#### 4.3.5 Sampling

Finally, one can draw samples from the GMMs to verify the clustering. To do so, we first chose one distribution (or cluster *c*) from the mixture, our choice weighted by the probability *p*_*c*_. It is then possible to generate the vector of lengths *l*_*a*_ or terminal points *t*_*a*_ by drawing a sample from the chosen distribution:

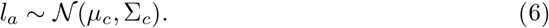

We added a post-processing step to all samples, which removes values sampled in regions not observed in the biological input data.

We used the sampling here as an indicator of the clustering accuracy only. However, one could use the sampling to synthesize directly axons with the sampled lengths, for instance by choosing tufts to add up to all lengths in each targeted region.

### 4.4 Synthesizing in the mouse brain

We now present how we used the axonal projections analysis presented in section 4.3 as input data for the synthesis algorithm 4.2 to synthesize LRAs in the mouse brain.

#### Initial morphologies synthesis

First of all, we synthesized neuronal morphologies made of somata, dendrites, and grafted reconstructed *local* axons in the isocortex, with the same methodology as in [7, 44]. These local axons were copied from previous experimental reconstructions and grafted to the synthesized cells. Since mostly pyramidal cells of the mouse cortex project to distal regions [45], we filtered pyramidal cells of the brain region *s*_*a*_ and would replace their local axon with a synthesized LRA. We synthesized LRAs for all pyramidal cells in the regions for which we synthesize, but that can be changed to a provided portion of them.

#### Source points

The somata of the filtered pyramidal cells were used as the source points of the synthesis algorithm. We assigned a GMM cluster for each source point by randomly picking a cluster *c* with probability *p*_*c*_ from the clusters *C* of source *s*_*a*_.

#### Target points

We computed the probability of an axon to target a brain region *b* as the number of axons targeting *b*, divided by the total number of axons in the cluster. For a picked target brain region, a number 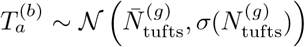 of target points were randomly placed inside *b*. All the targets of axon *a* were then connected with a better edge weight for targets inside fiber tracts to form the trunk. In this way, the trunk would preferentially follow the fiber tracts.

#### Tufts selection

Finally, tufts were selected for each target point with probability computed based on the representativity scores of their group *g*, and then synthesized with the NeuroTS software [7].

#### Creating connections

Once all the morphologies were fully synthesized, we made axo-dendritic connections as described in [24]. In a few words, touches were detected based on the physical proximity of neurite branches and filtered based on a minimum inter-bouton interval. Touches were then pruned based on physiological synapse density and converted into synapses. Connections and connectivity matrices were analyzed using the ConnectomeUtilities software [46].

## 5 Acknowledgements

This study was supported by funding to the Blue Brain Project, a research center of the Ećole polytechnique fédérale de Lausanne (EPFL), from the Swiss government’s ETH Board of the Swiss Federal Institutes of Technology. We further thank Cyrille Favreau, Fabien Petitjean and the Scientific Visualization team for developing the visualization tools that helped produce the rendering of circuits within the mouse brain and the Data Knowledge Engineering team for helping manage the morphologies datasets.

## 6 Data availability

Accompanying data can be found at https://doi.org/10.5281/zenodo.13790069 and scripts for data analysis at https://github.com/Remy2506/axon_projection_figures.git.

## 7 Supplementary Materials

### 7.1 Acronyms

A list of brain regions acronyms used can be found in table S1.

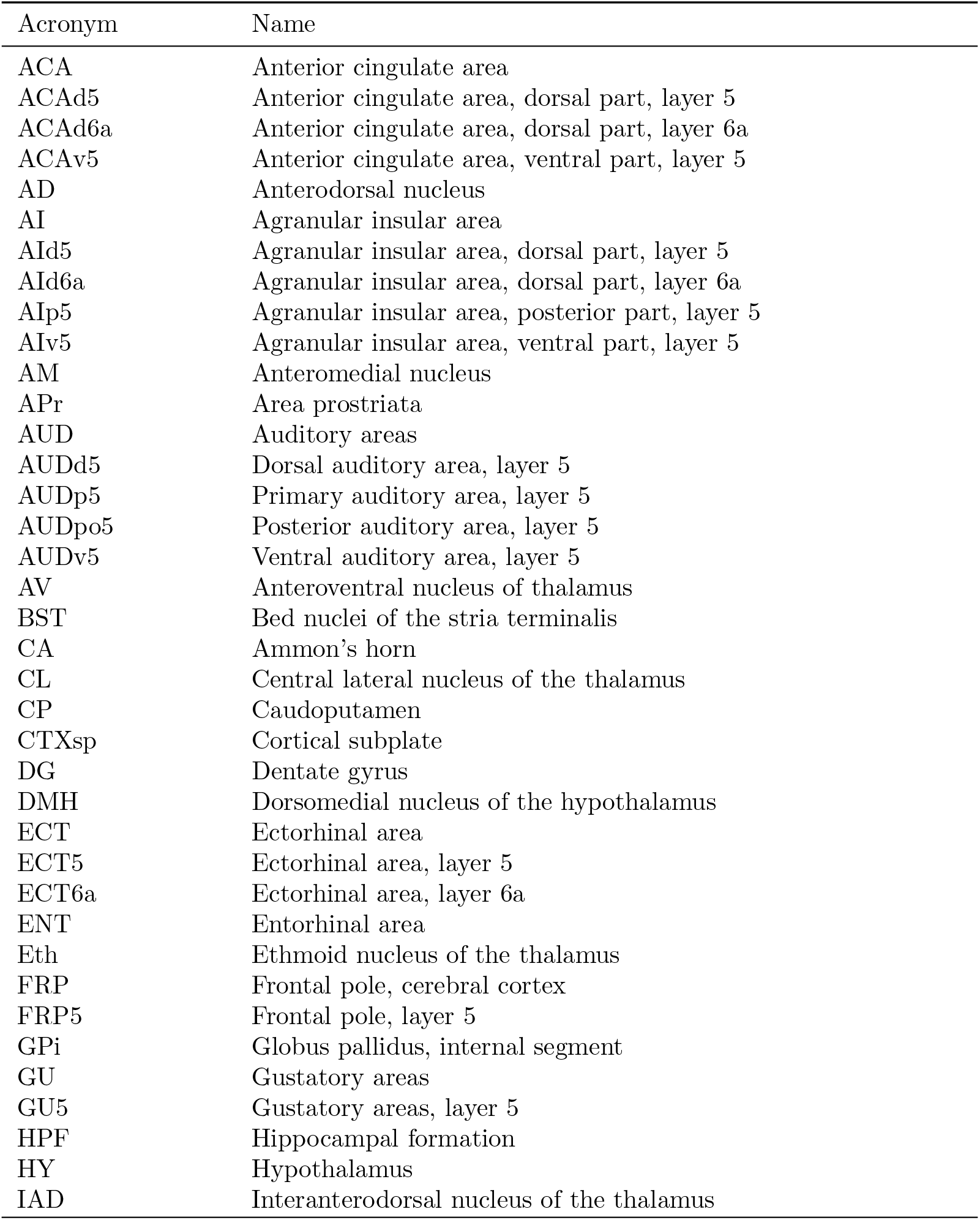
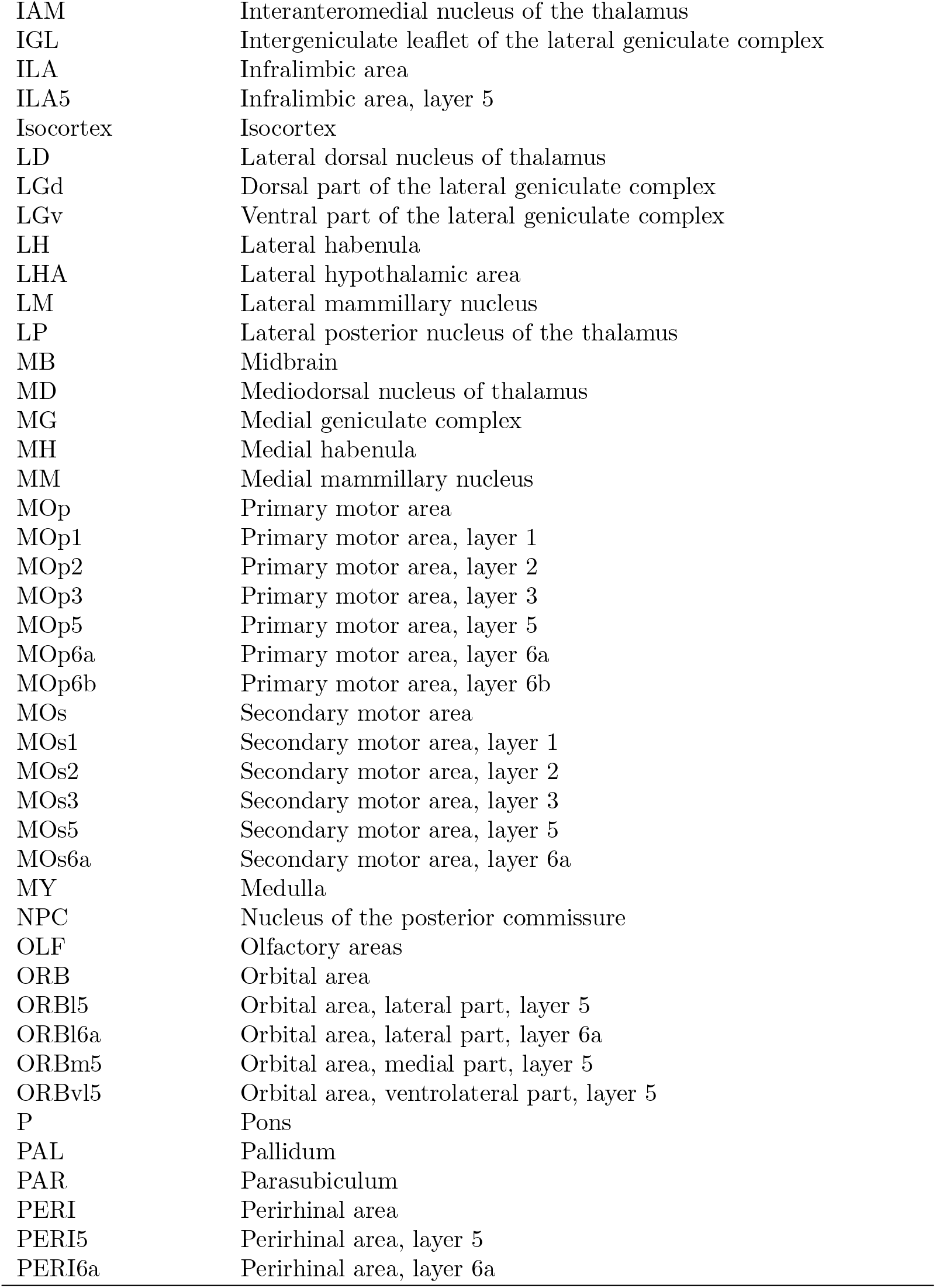
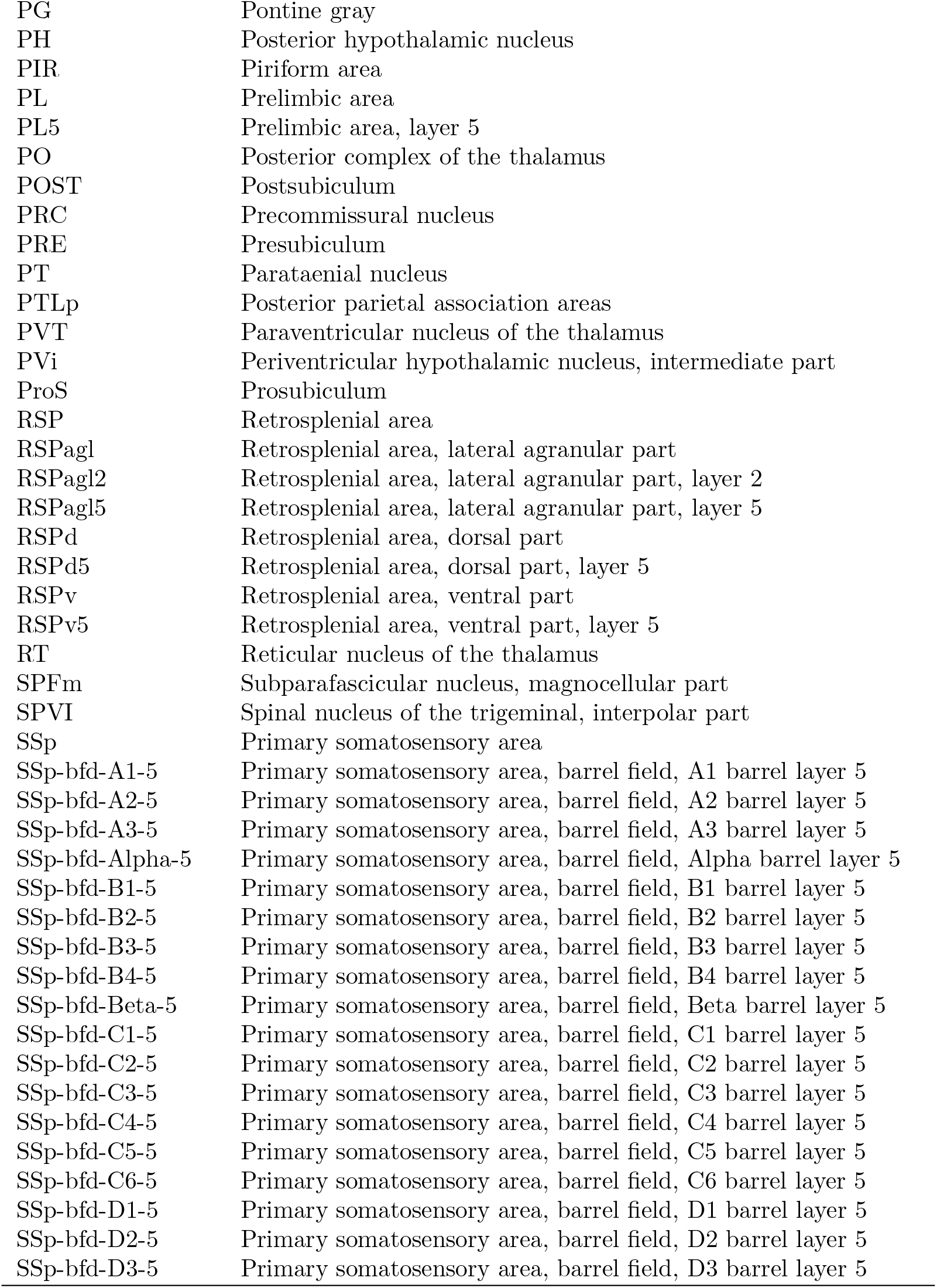
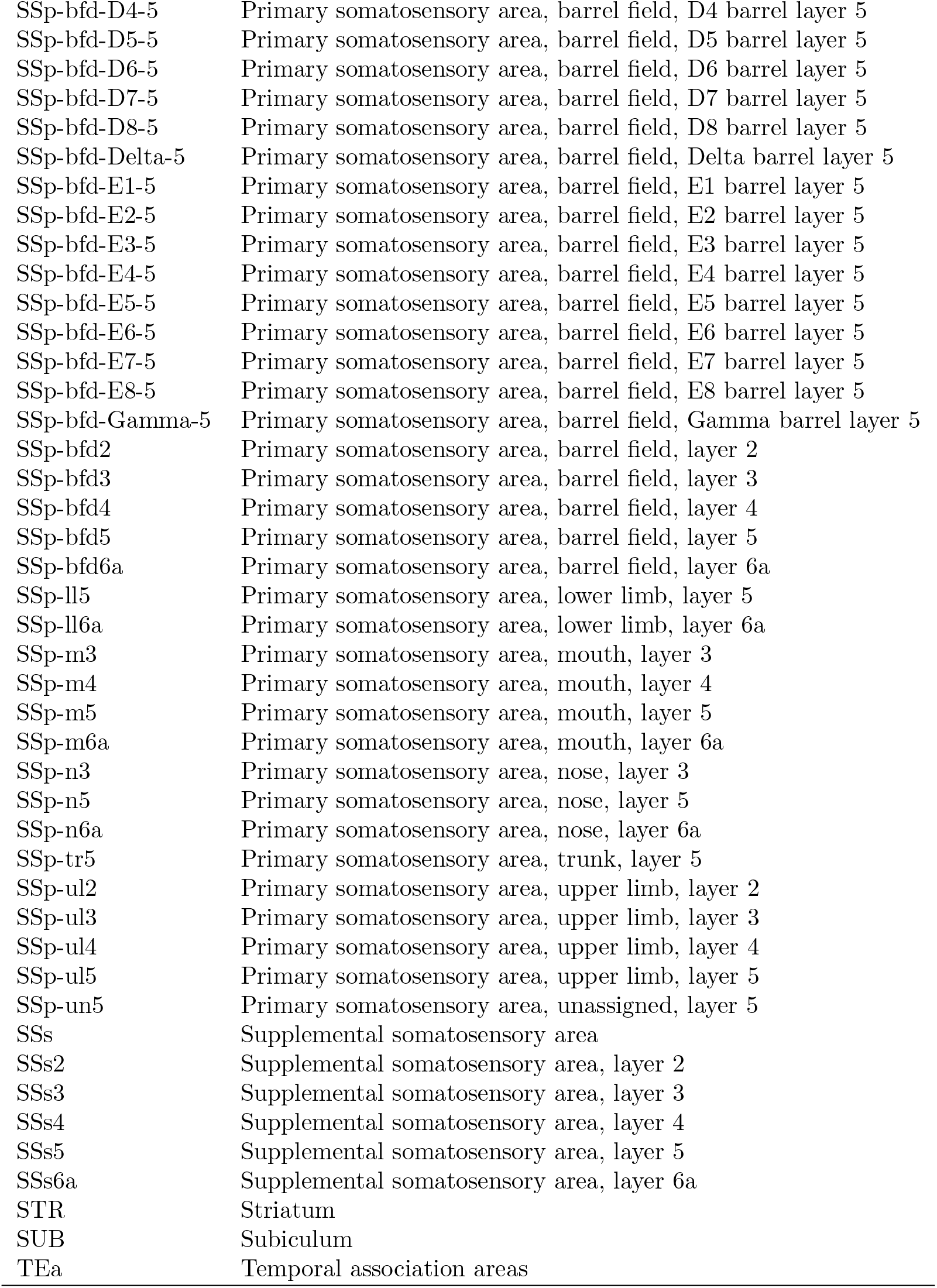
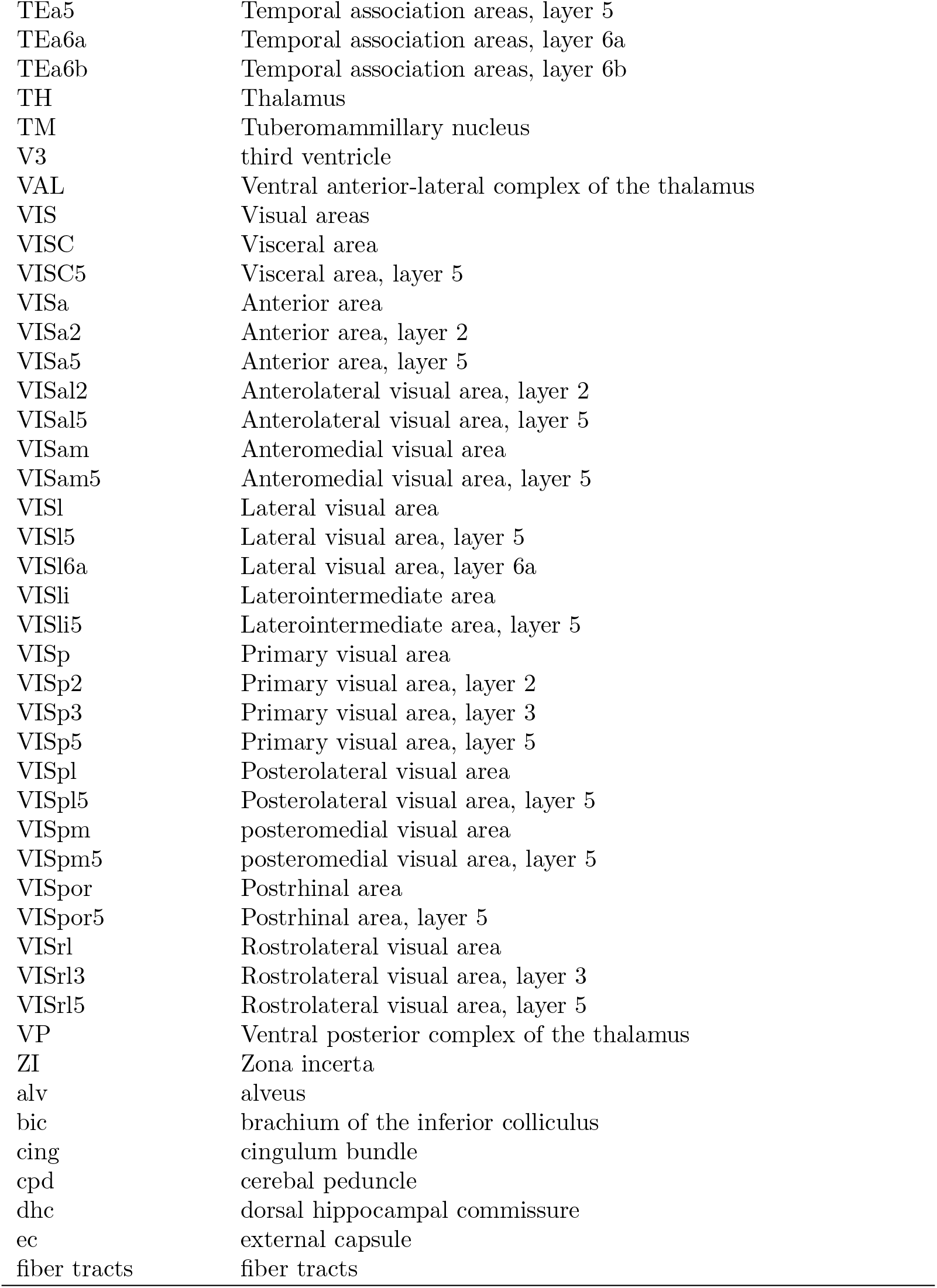

**Fig. S1.**
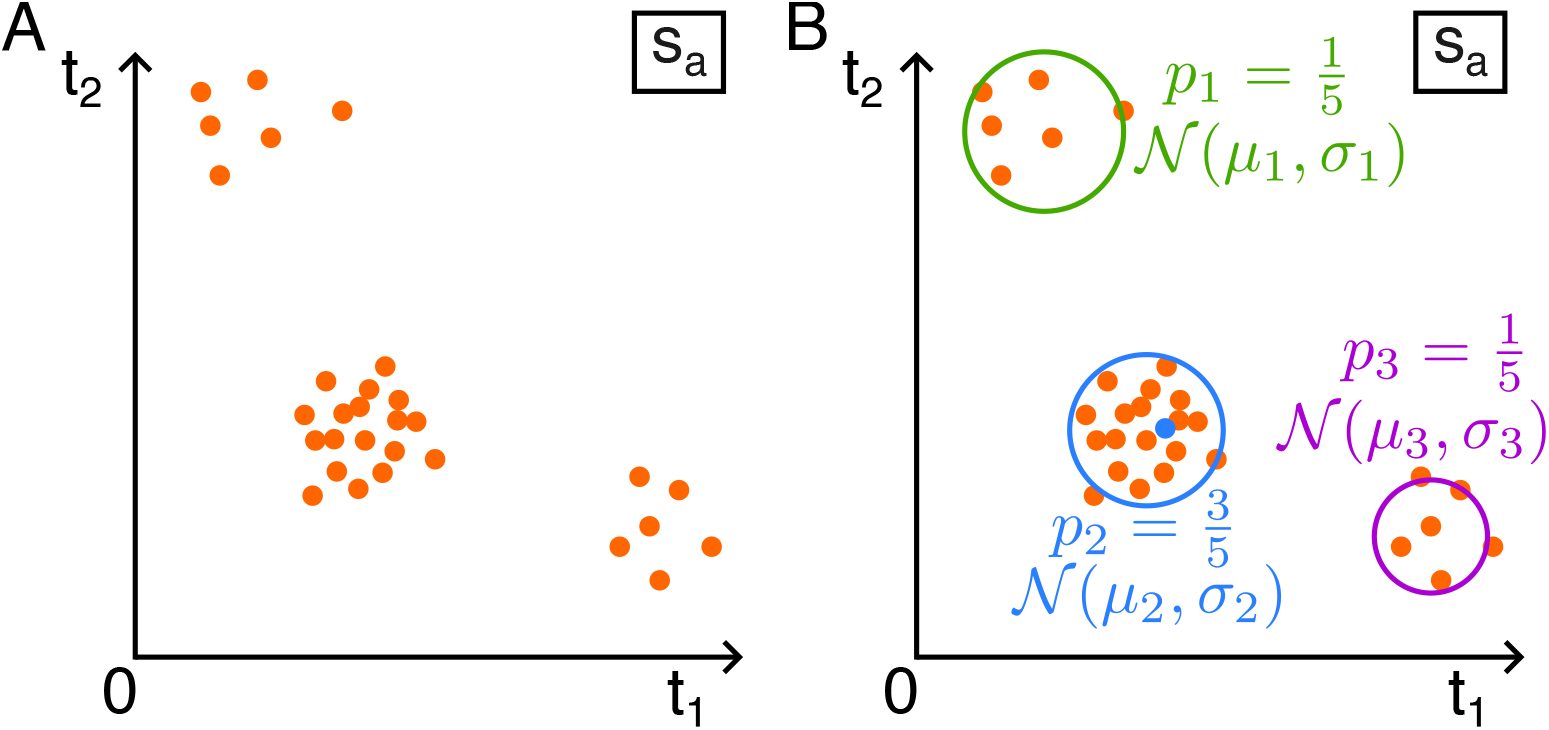
Schematic of the principle of Gaussian Mixture clustering, with axons starting in source region *s*_*a*_ and targeting only two target regions *t*_1_ and *t*_2_ (**A**). Each point is the feature vector value of an axon, e.g., number of terminals or axonal path length in *t*_1_ and *t*_2_. The clusters *C* are found using the EM algorithm 7.2.1 (**B**).

**Table S1.**
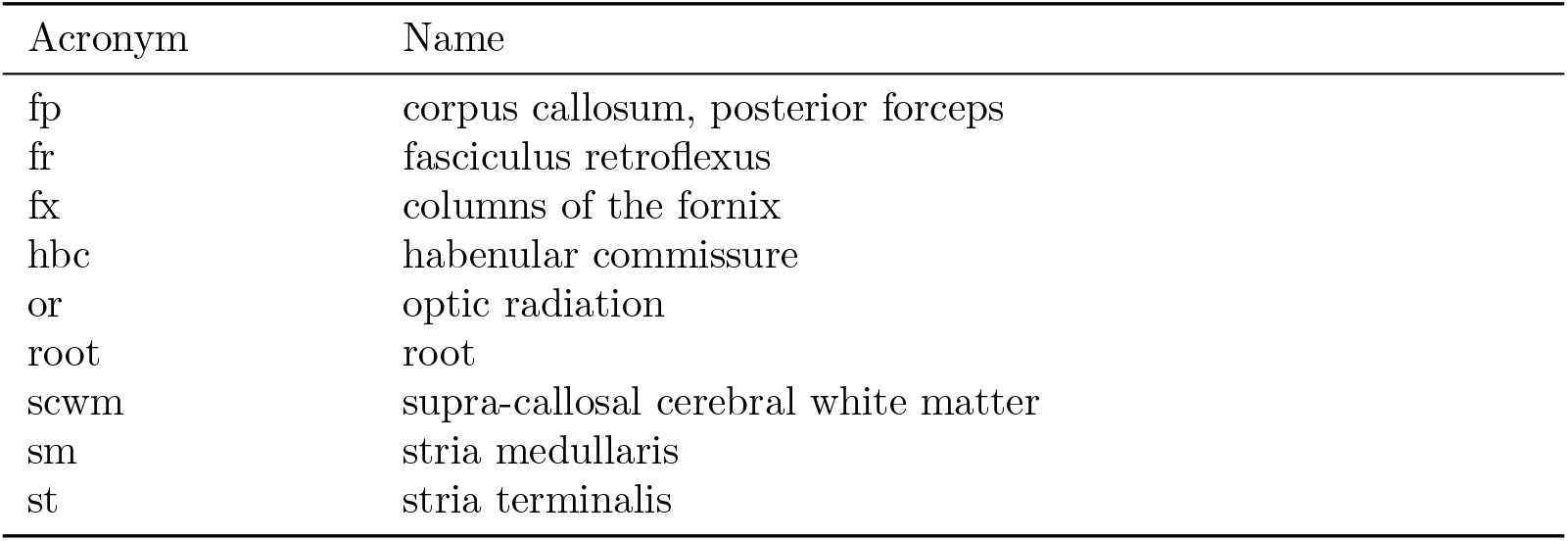
List of region acronyms used and their corresponding names.

### 7.2 Clustering formalism

A schematic illustration of the Gaussian Mixture clustering process can be found in Fig. S1. In Fig. S1A, we imagine a case where axons start from source region *s*_*a*_ and target only two target regions *t*_1_ and *t*_2_. Each point on the plot is the feature vector value of an axon, e.g., number of terminals or axonal path length in regions *t*_1_ and *t*_2_. Each cluster *c* is defined by a probability to belong to it *p*_*c*_, and a Gaussian distribution with mean feature values *µ*_*c*_ and standard deviation *σ*_*c*_, see Fig. S1B. The clusters *C* are found using the EM algorithm 7.2.1. The probability of falling into a cluster *p*_*c*_ is proportional to the number of axons in that cluster. Finally, one can sample feature vector values from a cluster using its normal distribution.

#### 7.2.1 EM algorithm

The EM-algorithm works in two steps.

##### Initialization

We start by initializing the parameters of the mixture models *θ*_0_. This can be done in the following straightforward way:

- For *µ*_*c*_, pick *C* random points among the data.
- For Σ_*c*_, compute the covariance matrix of the data for each cluster with *µ*_*c*_ from the previous step.
- Initialize 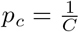 uniformly.

##### E step

In the E-step, we find the probability of *a* to belong to cluster *c* (also called posterior) by computing eq. (3), given a set of estimated parameters 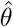. In doing so, we apply a soft clustering to our data, i.e., we give a probability of belonging to each class for each neuron of a source region. (Hard clustering means saying to which cluster belongs the neurons)

##### M step

In the M-step, given the assigned clustering probabilities, we compute a proxy 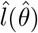 for the log-likelihood of the data and find the new estimated parameters 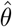 that maximize it. Let this proxy be:

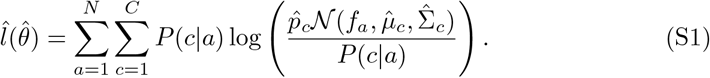

Taking the derivatives of eq. (S1) with respect to 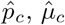 and 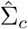 and setting them equal to zero, we get the new estimates to be used in the next E-step :

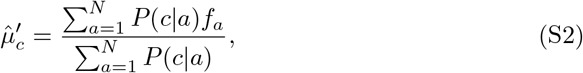

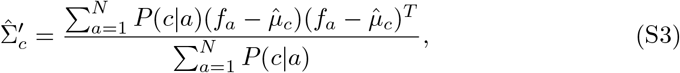

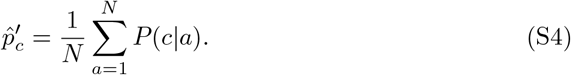

##### Termination

We iterate the E- and M-steps until we reach a (local) maximum for 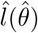, which is guaranteed [47]. We can set the termination condition to be something like 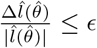, with *ϵ* ∈ ℝ small.

#### 7.2.2 Number of clusters

A limitation of the GMM clustering is that it needs to know *a priori* the number of clusters *C* to expect for each source region. One way to estimate *C* is to compute the likelihood of the data for different values of *C*. Of course, the likelihood will be maximal in the limit where *C* = *N*, but that would not be what we are looking for. A method to balance the log-likelihood by the number of clusters is to take the minimal Bayesian Information Criterion (BIC) for the model, defined as [48]:

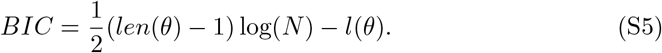

(*len*(*θ*) − 1) is the number of free parameters of the model; in our case it is *C* − 1 + *C × B* + *C*, because *θ* = {*p*_1_, …, *p*_*C*_, *µ*_1_, …, *µ*_*C*_, Σ_1_, …, Σ_*C*_} and :

- 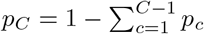, thus we get *C* − 1 from the probabilities,
- The means *µ*_*c*_ ∈ ℝ^*B*^ are of the dimension of the data, thus *C × B* from the means.
- We get *C* free parameters for the variances because the elements of Σ_*c*_ are products from *C* values *σ*_*c*_ ∈ ℝ.

#### 7.2.3 Limitations and possible improvements

Here, we give a non-exhaustive list of limitations and possible improvements to the Gaussian clustering method presented in this work.

- The local maxima reached in the GMMs with EM are sensitive to the initial parameters *θ*_0_. A common approach is to use the K-means algorithm to initialize *θ*_0_, which we used here. However, other methods can be used (e.g., random initialization). We did not study the impact of initialization.
- If the local maxima given by EM are not good enough, we can use other methods to find *θ*, such as the method of moments [49].
- It would be better to know *a priori* the expected number of clusters for each source. Here, we use tools like the Bayesian Information Criterion to choose a balanced number of clusters with respect to the number of parameters. We could think of choosing the number of clusters with another clustering method (e.g., hierarchical clustering made in [29]). But we don’t give it too much importance in this work because we focus on reproducing axon targeting, not on giving an ontological classification.
- We did not consider morphologynor transcriptomic-types of the morphologies. This could be added to the feature vector for clustering.

### 7.3 Tufts representativity score

We computed a *representativity score R* for each tuft *τ* of a given group *g*, defined by GMM cluster and the target region. This score measures how much a set of *M* morphometrical features of a tuft *τ* are close to those of tufts *T* (*g*) of its group.

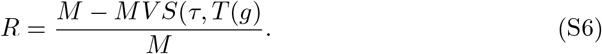

**Fig. S2.**
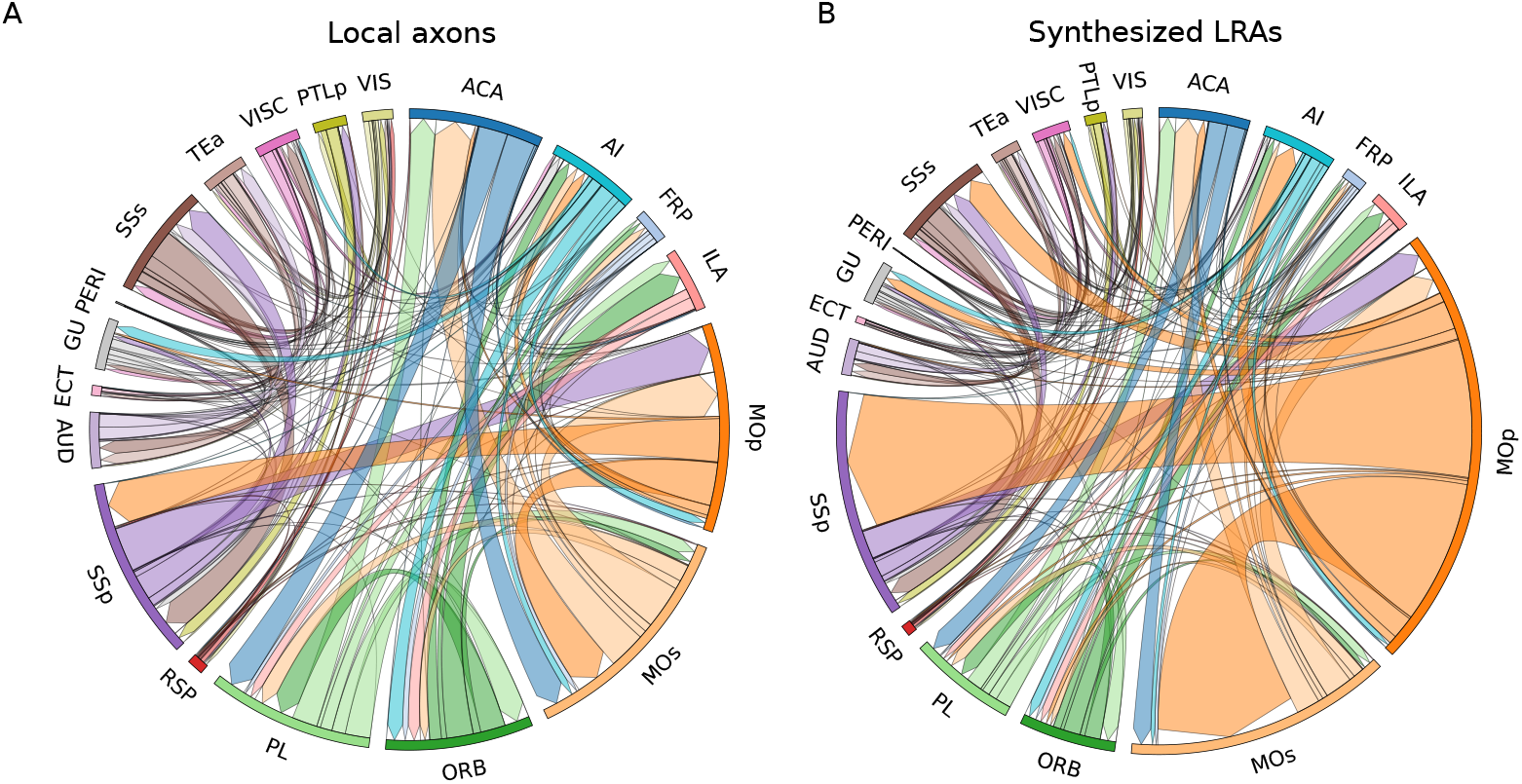
Outgoing connections of axons in the isocortex. **A**: Only local axons are used. **B**: LRAs are synthesized for MOp5 axons.

We used `NeuroM` [43] to compute the following set of morphometrics: section lengths, remote bifurcation angles, number of sections per neurite, terminal path lengths, section terminal branch orders, section path distances, section terminal lengths, section terminal radial distances.

We used this score as a pick probability for the tufts synthesis.

### 7.4 Supplementary figures

We show in Fig. S2 the outgoing connectivity of the synthesized MOp5 axons presented in Fig. 4C and D.

In Fig. S3, we show additional views of the reconstructed and synthesized LRAs from the MOp5 region, with a set of highlighted regions: the caudoputamen (CP), the primary and secondary motor areas (MOp, MOs), the pontine gray region (PG), the spinal nucleus of the trigeminal interpolar part (SPVI), and the primary and secondary somatosensory areas (SSp, SSs).

In Fig. S4, we compared the distributions of neurite lengths between reconstructed local axons and LRAs. Fig. S4A shows the axons of the MOp5 region, and S4B the whole sets of axons.

### 7.5 Biological LRAs in the isocortex

**Table S2.**
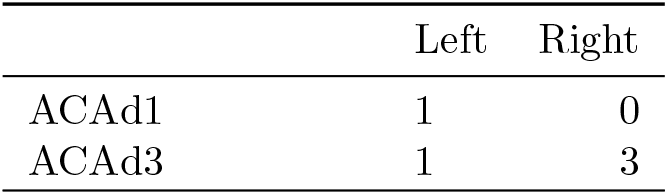
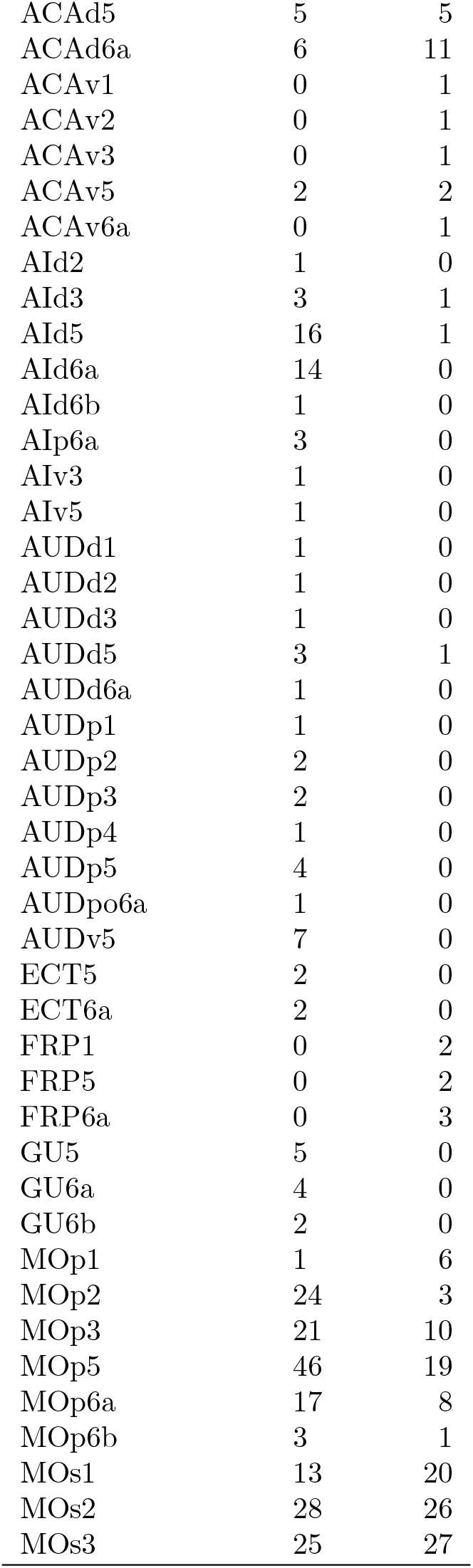
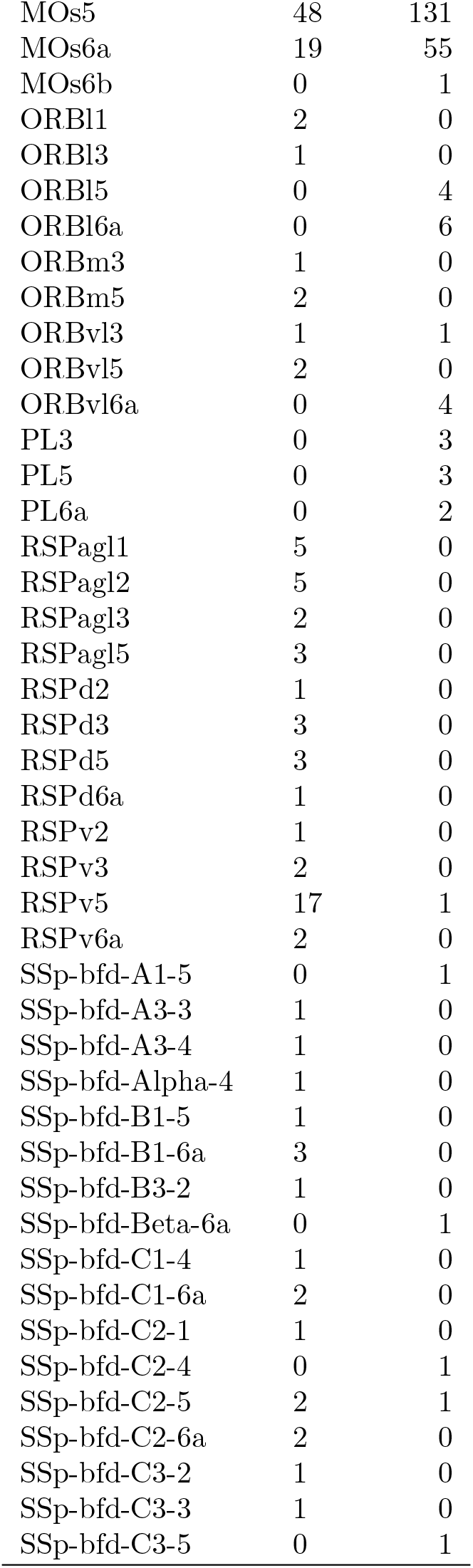
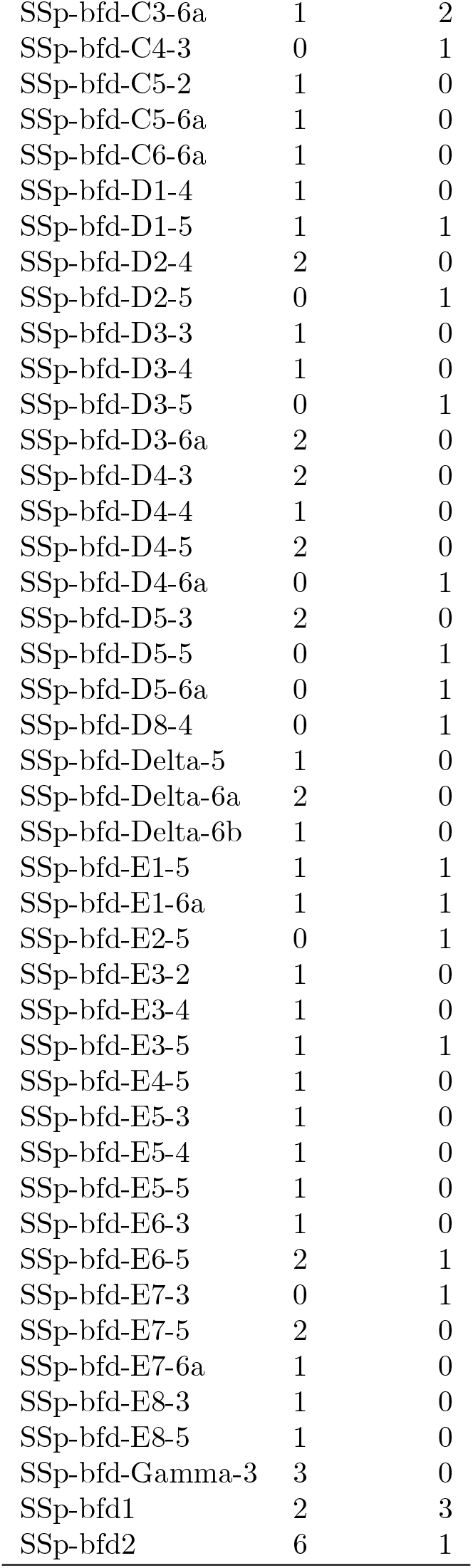
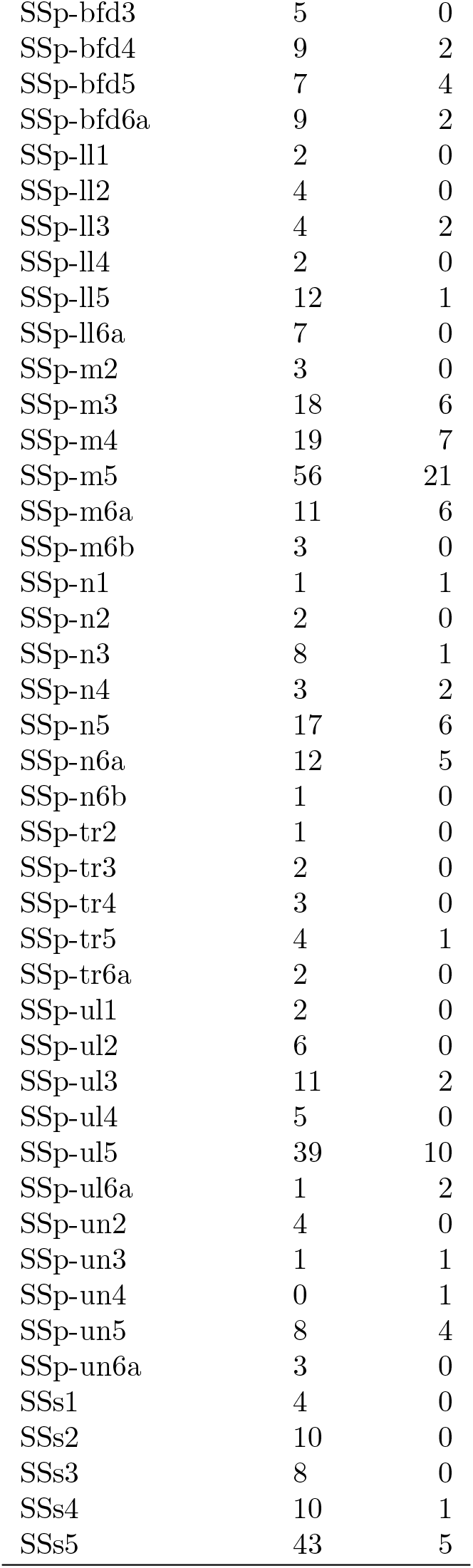
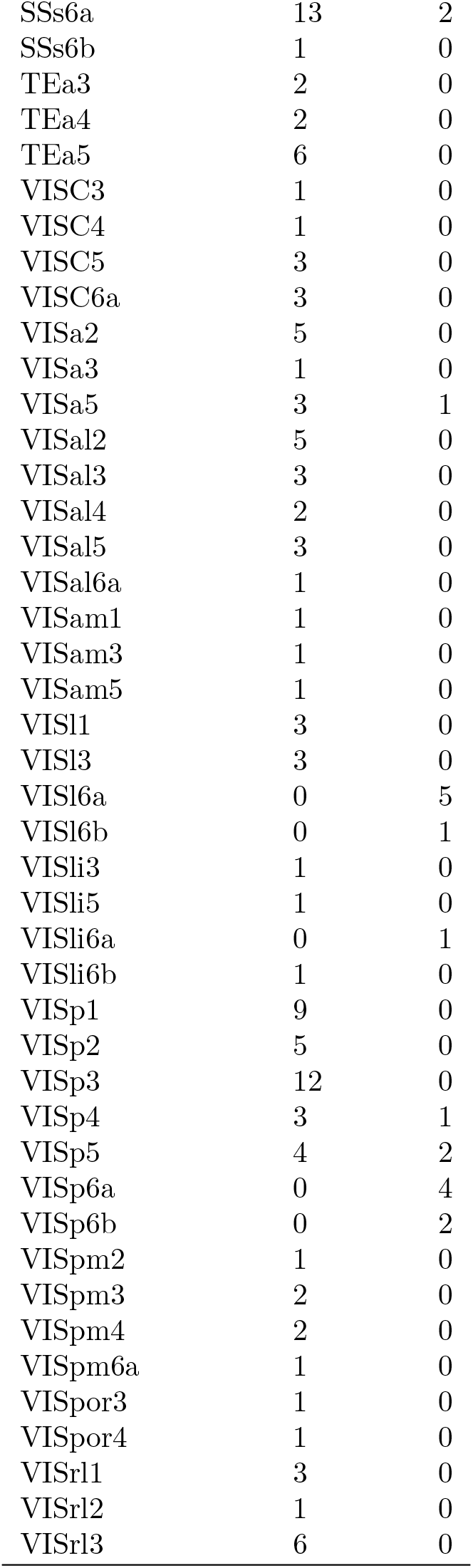
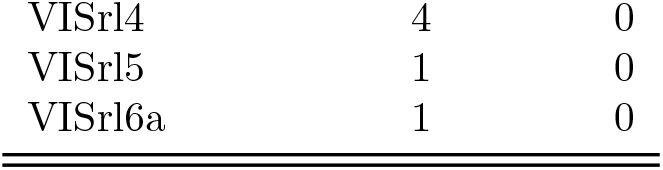
Number of axons in the biological input data originating from the isocortex, by hemisphere.

**Fig. S3.**
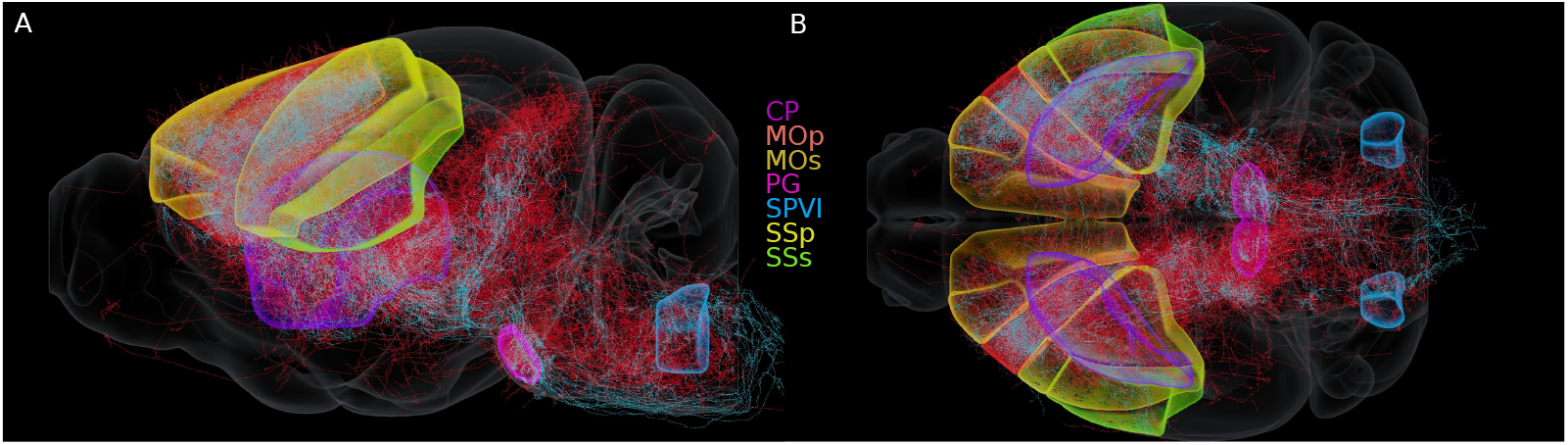
The 65 reconstructed axons and 170 of the synthesized axons in the mouse brain atlas. Some regions of interest, inside and outside of the isocortex, are highlighted: caudoputamen (CP), the primary and secondary motor areas (MOp, MOs), the pontine gray region (PG), the spinal nucleus of the trigeminal interpolar part (SPVI), and the primary and secondary somatosensory areas (SSp, SSs). **A**: Lateral view. **B**: Top view.

**Fig. S4.**
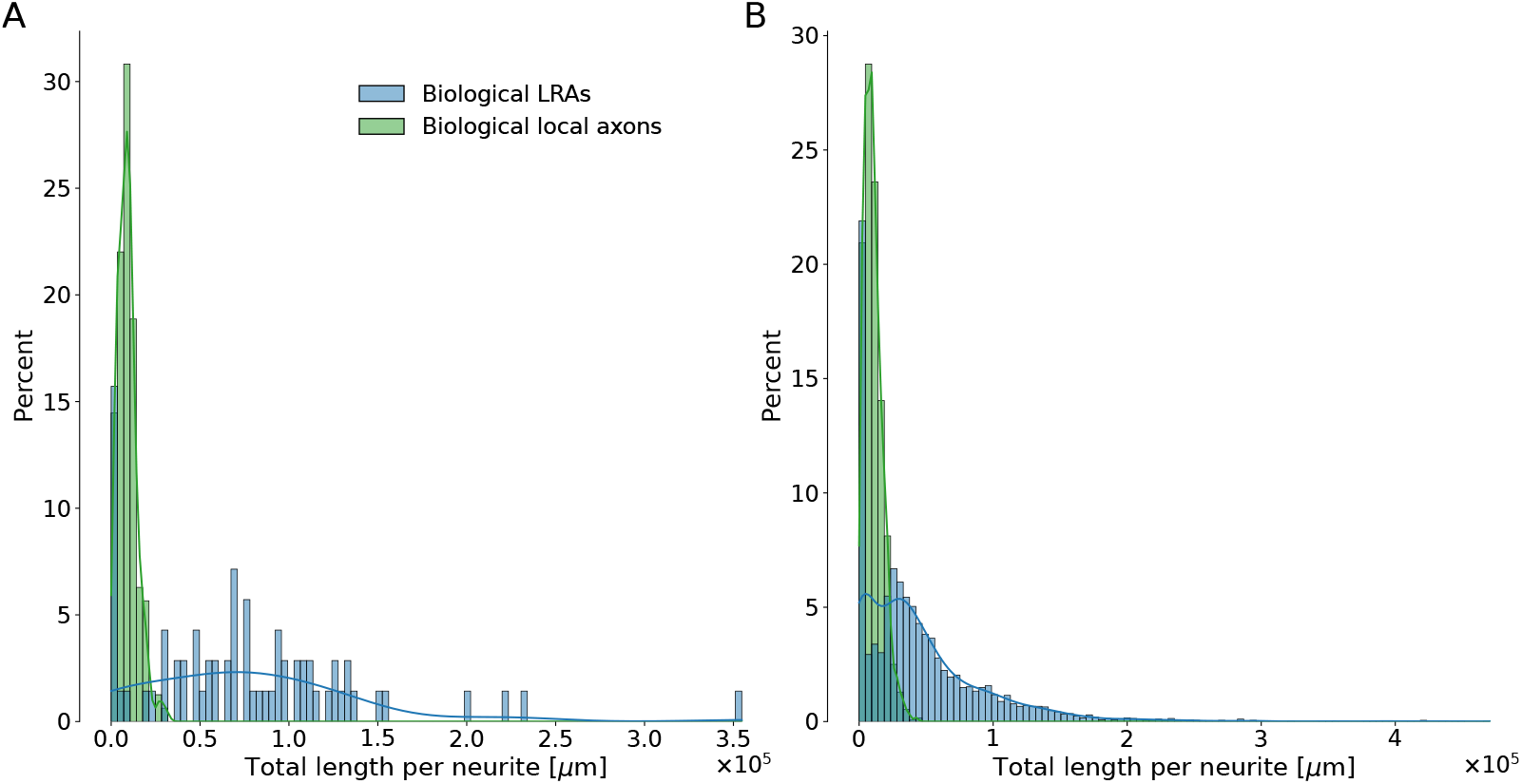
Distribution of total lengths per neurite for biological local axons previously used in [7] (green) and reconstructions of LRAs from the dataset used in the present work (blue). **A**: Considering only axons used in the MOp5 region. **B**: Considering all axons from both datasets.

## Notes

https://doi.org/10.5281/zenodo.13790069

### Competing Interest Statement

The authors have declared no competing interest.

### Summary of Updates

Correction of minor typographical errors. Exhaustive list: - Corrected legend of Figs 2 and 6 - Reformulated a few sentences in discussion - Added DOI of axon-synthesis article - Typo '?' in Methods - Input morphologies

## References

[1] DeFelipe, J. The anatomical problem posed by brain complexity and size: a potential solution. Frontiers in Neuroanatomy 9 (2015). URL https://www.frontiersin.org/journals/neuroanatomy/articles/10.3389/fnana.2015.00104/full. Publisher: Frontiers.

[2] Zubler, F. & Douglas, R. A framework for modeling the growth and development of neurons and networks. Frontiers in Computational Neuroscience 3 (2009).

[3] Cuntz, H., Forstner, F., Borst, A. & Häusser, M. One Rule to Grow Them All: A General Theory of Neuronal Branching and Its Practical Application. PLOS Computational Biology 6, e1000877 (2010). URL https://journals.plos.org/ploscompbiol/article?id=10.1371/journal.pcbi.1000877. Publisher: Public Library of Science.

[4] Luczak, A. Measuring Neuronal Branching Patterns Using Model-Based Approach. Frontiers in Computational Neuroscience 4 (2010). URL https://www.frontiersin.org/journals/computational-neuroscience/articles/10.3389/fncom.2010.00135/full. Publisher: Frontiers.

[5] Ascoli, G. et al. Computer generation and quantitative morphometric analysis of virtual neurons. Anatomy and Embryology 204, 283–301 (2001).

[6] Koene, R. A. et al. NETMORPH: A Framework for the Stochastic Generation of Large Scale Neuronal Networks With Realistic Neuron Morphologies. Neuroinformatics 7, 195–210 (2009). URL 10.1007/s12021-009-9052-3.

[7] Kanari, L. et al. Computational synthesis of cortical dendritic morphologies. Cell Reports 39, 110586 (2022). URL https://www.sciencedirect.com/science/article/pii/S2211124722003308.

[8] Kanari, L. et al. A Topological Representation of Branching Neuronal Morphologies. Neuroinformatics 16, 3–13 (2018). URL 10.1007/s12021-017-9341-1.

[9] Kanari, L. et al. Objective Morphological Classification of Neocortical Pyramidal Cells. Cerebral Cortex 29, 1719–1735 (2019). URL 10.1093/cercor/bhy339.

[10] Zisis, E. et al. Digital Reconstruction of the Neuro-Glia-Vascular Architecture. Cerebral Cortex (New York, NY) 31, 5686–5703 (2021). URL https://www.ncbi.nlm.nih.gov/pmc/articles/PMC8568010/.

[11] McAllister, A. K. Cellular and Molecular Mechanisms of Dendrite Growth. Cerebral Cortex 10, 963–973 (2000). URL 10.1093/cercor/10.10.963.

[12] Sugimura, K., Shimono, K., Uemura, T. & Mochizuki, A. Self-organizing Mechanism for Development of Space-filling Neuronal Dendrites. PLOS Computational Biology 3, e212 (2007). URL https://journals.plos.org/ploscompbiol/article?id=10.1371/journal.pcbi.0030212. Publisher: Public Library of Science.

[13] Kirchner, J. H., Euler, L. & Gjorgjieva, J. Dendritic growth and synaptic organization from activity-independent cues and local activity-dependent plasticity. eLife 12 (2023). URL https://elifesciences.org/reviewed-preprints/87527. Publisher: eLife Sciences Publications Limited.

[14] Gibson, D. A. & Ma, L. Developmental regulation of axon branching in the vertebrate nervous system. Development 138, 183–195 (2011). URL 10.1242/dev.046441.

[15] Vanherpe, L., Kanari, L., Atenekeng, G., Palacios, J. & Shillcock, J. Framework for efficient synthesis of spatially embedded morphologies. Phys. Rev. E 94, 023315 (2016). URL https://link.aps.org/doi/10.1103/PhysRevE.94.023315.

[16] Costa, R. P. Computational model of axon guidance (2015). URL http://arxiv.org/abs/1508.01537. ArXiv:1508.01537 [q-bio].

[17] Budd, J. M. L. et al. Neocortical Axon Arbors Trade-off Material and Conduction Delay Conservation. PLOS Computational Biology 6, e1000711 (2010). URL https://journals.plos.org/ploscompbiol/article?id=10.1371/journal.pcbi.1000711. Publisher: Public Library of Science.

[18] Callaghan, R., Alexander, D. C., Palombo, M. & Zhang, H. ConFiG: Contextual Fibre Growth to generate realistic axonal packing for diffusion MRI simulation. NeuroImage 220, 117107 (2020). URL https://www.sciencedirect.com/science/article/pii/S1053811920305930.

[19] Bingham, C. S. et al. ROOTS: An Algorithm to Generate Biologically Realistic Cortical Axons and an Application to Electroceutical Modeling. Frontiers in Computational Neuroscience 14 (2020). URL https://www.frontiersin.org/journals/computational-neuroscience/articles/10.3389/fncom.2020.00013/full. Publisher: Frontiers.

[20] Ascoli, G. et al. Petilla terminology: Nomenclature of features of GABAergic interneurons of the cerebral cortex. Nature Reviews Neuroscience 9, 557–568 (2008).

[21] Yamahachi, H., Marik, S. A., McManus, J. N. J., Denk, W. & Gilbert, C. D. Rapid axonal sprouting and pruning accompany functional reorganization in primary visual cortex. Neuron 64, 719–729 (2009).

[22] Economo, M. N., Winnubst, J., Bas, E., Ferreira, T. A. & Chandrashekar, J. Single-neuron axonal reconstruction: The search for a wiring diagram of the brain. Journal of Comparative Neurology 527, 2190– 2199 (2019). URL https://onlinelibrary.wiley.com/doi/abs/10.1002/cne.24674.eprint: https://onlinelibrary.wiley.com/doi/pdf/10.1002/cne.24674.

[23] Reimann, M. W. et al. A null model of the mouse whole-neocortex microconnectome. Nature Communications 10, 3903 (2019). URL https://www.nature.com/articles/s41467-019-11630-x.

[24] Markram, H. et al. Reconstruction and Simulation of Neocortical Microcircuitry. Cell 163, 456–492 (2015). URL https://linkinghub.elsevier.com/retrieve/pii/S0092867415011915.

[25] Berchet, A., Petkantchin, R., Markram, H. & Kanari, L. Computational generation of long-range axonal morphologies (2024). URL https://www.biorxiv.org/content/10.1101/2024.10.16.618695v1. Pages: 2024.10.16.618695 Section: New Results.

[26] Winnubst, J. et al. Reconstruction of 1,000 projection neurons reveals new cell types and organization of long-range connectivity in the mouse brain. Cell 179, 268–281.e13 (2019). URL https://www.ncbi.nlm.nih.gov/pmc/articles/PMC6754285/.

[27] Peng, H. et al. Morphological diversity of single neurons in molecularly defined cell types. Nature 598, 174–181 (2021). URL https://www.nature.com/articles/s41586-021-03941-1. Number: 7879 Publisher: Nature Publishing Group.

[28] Jiang, S. et al. Anatomically revealed morphological patterns of pyramidal neurons in layer 5 of the motor cortex. Scientific Reports 10, 7916 (2020). URL https://www.nature.com/articles/s41598-020-64665-2. Publisher: Nature Publishing Group.

[29] Wheeler, D. W., Banduri, S., Sankararaman, S., Vinay, S. & Ascoli, G. A. Unsupervised classification of brain-wide axons reveals neuronal projection blueprint. Research Square rs.3.rs–3044664 (2023). URL https://www.ncbi.nlm.nih.gov/pmc/articles/PMC10350180/.

[30] Bolaños-Puchet, S. et al. Enhancement of brain atlases with laminar coordinate systems: Flatmaps and barrel column annotations. Imaging Neuroscience 2, 1–20 (2024). URL 10.1162/imaga00209.

[31] Nachbaur, D. et al. BlueBrain/Brayns: 3.6.0 Release (2024). URL https://zenodo.org/records/10694081.

[32] Cadwell, C. R. et al. Electrophysiological, transcriptomic and morphologic profiling of single neurons using patch-seq. Nature Biotechnology 34, 199–203 (2015). URL https://api.semanticscholar.org/CorpusID:344546.

[33] Scala, F. et al. Phenotypic variation of transcriptomic cell types in mouse motor cortex. Nature 598, 144 – 150 (2020). URL https://api.semanticscholar.org/CorpusID:226851177.

[34] Gouwens, N. W. et al. Integrated morphoelectric and transcriptomic classification of cortical gabaergic cells. Cell 183, 935–953.e19 (2020). URL https://api.semanticscholar.org/CorpusID:226308670.

[35] Markram, H. The Blue Brain Project. Nature Reviews Neuroscience 7, 153–160 (2006). URL https://www.nature.com/articles/nrn1848. Number: 2 Publisher: Nature Publishing Group.

[36] Palombo, M. et al. Sandi: A compartment-based model for non-invasive apparent soma and neurite imaging by diffusion mri. Neuroimage 215 (2019). URL https://api.semanticscholar.org/CorpusID:195820546.

[37] Wang, Q. et al. The Allen Mouse Brain Common Coordinate Framework: A 3D Reference Atlas. Cell 181, 936–953.e20 (2020).

[38] Hegde, C., Indyk, P. & Schmidt, L. A Nearly-Linear Time Framework for Graph-Structured Sparsity .

[39] Berchet, A., Arnaudon, A. & alex4200. BlueBrain/morphology-workflows: 0.11.0 (2024). URL https://zenodo.org/records/11072355.

[40] Ikotun, A. M., Ezugwu, A. E., Abualigah, L., Abuhaija, B. & Heming, J. K-means clustering algorithms: A comprehensive review, variants analysis, and advances in the era of big data. Information Sciences 622, 178–210 (2023). URL https://www.sciencedirect.com/science/article/pii/S0020025522014633.

[41] Reynolds, D. in Gaussian Mixture Models (eds Li, S.Z. & Jain, A.) Encyclopedia of Biometrics 659–663 (Springer US, Boston, MA, 2009). URL 10.1007/978-0-387-73003-5196.

[42] Ari, C., Aksoy, S. & Arikan, O. Maximum likelihood estimation of Gaussian mixture models using stochastic search. Pattern Recognition 45, 2804–2816 (2012). URL https://www.sciencedirect.com/science/article/pii/S0031320312000167.

[43] Arnaudon, A. et al. NeuroM (2024). URL https://zenodo.org/records/10630119.

[44] Arnaudon, A. et al. Controlling morpho-electrophysiological variability of neurons with detailed biophysical models (2023). URL https://www.biorxiv.org/content/10.1101/2023.04.06.535923v1. Pages: 2023.04.06.535923 Section: New Results.

[45] Gerfen, C. R., Economo, M. N. & Chandrashekar, J. Long Distance Projections of Cortical Pyramidal Neurons. Journal of neuroscience research 96, 1467–1475 (2018). URL https://www.ncbi.nlm.nih.gov/pmc/articles/PMC5429214/.

[46] Reimann, M., Ecker, A., Sato, A., Kurban, K. & Lupton, O. BlueBrain/ConnectomeUtilities: New features and examples: Reciprocal views (2023). URL https://zenodo.org/records/10059227.

[47] Wu, C. F. J. On the Convergence Properties of the EM Algorithm. The Annals of Statistics 11, 95–103 (1983). URL https://projecteuclid.org/journals/annals-of-statistics/volume-11/issue-1/On-the-Convergence-Properties-of-the-EM-Algorithm/10.1214/aos/1176346060.full. Publisher: Institute of Mathematical Statistics.

[48] Neath, A. A. & Cavanaugh, J. E. The Bayesian information criterion: background, derivation, and applications. WIREs Computational Statistics 4, 199–203 (2012). URL https://onlinelibrary.wiley.com/doi/abs/10.1002/wics.199. eprint: https://onlinelibrary.wiley.com/doi/pdf/10.1002/wics.199.

[49] Wu, Y. & Yang, P. Optimal estimation of Gaussian mixtures via denoised method of moments (2019). URL http://arxiv.org/abs/1807.07237. 1807.07237 [math, stat].

